# Mechanotransduction governs CD40 function and underlies X-linked Hyper IgM syndrome

**DOI:** 10.1101/2023.07.23.550231

**Authors:** Hyun-Kyu Choi, Stefano Travaglino, Matthias Münchhalfen, Richard Görg, Zhe Zhong, Jintian Lyu, David M. Reyes-Aguilar, Jürgen Wienands, Ankur Singh, Cheng Zhu

## Abstract

B cell maturation in germinal centers (GCs) depends on cognate interactions between the T and B cells. Upon interaction with CD40 ligand (CD40L) on T cells, CD40 delivers co-stimulatory signals alongside B cell antigen receptor (BCR) signaling to regulate affinity maturation and antibody class-switch during GC reaction. Mutations in CD40L disrupt interactions with CD40, which lead to abnormal antibody responses in immune deficiencies known as X-linked Hyper IgM syndrome (X-HIgM). Assuming that physical interactions between highly mobile T and B cells generate mechanical forces on CD40–CD40L bonds, we set out to study the B cell mechanobiology mediated by CD40–CD40L interaction. Using a suite of biophysical assays we find that CD40 forms catch bond with CD40L where the bond lasts longer at larger forces, B cells exert tension on CD40–CD40L bonds, and force enhances CD40 signaling and antibody class-switch. Significantly, X-HIgM CD40L mutations impair catch bond formation, suppress endogenous tension, and reduce force-enhanced CD40 signaling, leading to deficiencies in antibody class switch. Our findings highlight the critical role of mechanotransduction in CD40 function and provide insights into the molecular mechanisms underlying X-HIgM syndrome.

## Introduction

As a key costimulatory receptor on B cells, CD40 delivers complementary signals alongside B cell antigen receptor (BCR) signaling to prevent B cell silencing or deletion.^1–3^ Upon binding to the BCR, a thymus-dependent (TD) antigen is efficiently internalized by the B cell and the processed peptides are presented by major histocompatibility complex (pMHC) molecules on the cell surface. This TD antigen pMHC is tested by the T cell receptor (TCR) of follicular T helper (T_FH_) cells in a process known as linked recognition.^4,5^ If both the B and T_FH_ cells recognize the same TD antigen, the activated T_FH_ cell expresses a costimulatory ligand, CD40L (CD154)^6^, which binds CD40 on B cells. Such binding induces CD40 signaling to activate the non-canonical NF-kB pathway,^7^ promoting B cell activation and their terminal differentiation into antibody-secreting plasma cells.^8,9^ In addition to CD40 signaling, multiple cytokines regulate B cell response and differentiation, including B-cell activating factor (BAFF), produced by follicular dendritic cells (FDC)^8^, and IL-4 and IL-21, secreted by T_FH_ cells.^10^ B cells, FDCs, and T_FH_ cells form germinal centers (GCs) where B cells undergo proliferation and somatic hypermutation^11^ to generate B cell clones expressing a BCR with increased antigen affinity by a process known as affinity maturation. CD40 signaling is also critical for immunoglobulin (Ig) class switch recombination by which B cells can exchange the surface Ig isotype from IgM to other Ig classes, *i.e.*, IgG, IgE or IgA, which possess different immune effector functions and tissue distribution.^12,13^

Affinity maturation and Ig class-switch of GC B (GCB) cells require signaling induced by the cross T-B intercellular junctional interaction of CD40 with CD40L, which also generates a survival signal to prevent apoptosis induced by the BCR signal alone.^14^ B cells repeatedly migrate between the dark and light zones of GCs to take up cognate antigens from FDCs, and present the proteolytic peptide products *via* MHC class II to T_FH_ cells, thereby initiating the CD40–CD40L costimulatory signaling axis.^15–17^ In the absence of this interaction, B cells undergo apoptosis, allowing for the selection of B cell clones with a high-affinity BCR.^15,16^

Various genetic defects that hinder B cells to undergo class-switch and lead to abnormal antibody responses have been discovered, together making up a class of immune deficiencies known as X-linked Hyper IgM syndrome (X-HIgM).^18–23^ The most common defects leading to X-HIgM are mutations to the CD40L protein itself (X-HIgM type Ⅰ) that affect its expression, binding, or function,^18–20,24^ resulting in dysregulated signaling in B cells and their inability to undergo antibody class-switch.^19^ Other defects include the enzymes involved in class switch recombination, such as activation-induced cytidine deaminase^21^ uracil-DNA glycosylase.^22^ The inability of X-HIgM patients’ B cells to undergo class-switches from IgM to the IgG, IgA, or IgE isotypes renders them partially immunodeficient, unable to generate lasting immune memory to pathogens, and prone to recurring bacterial, fungal, or viral infections.^18,23,25^ Among the numerous possible mutations to CD40L, missense mutations are of high interest because they impact CD40–CD40L binding rather than trivially abrogating expression.^24^

CD40 is a category II Tumor Necrosis Factor receptor (TNFR). Unlike category I TNFRs, which can be activated by soluble TNF ligands (TNFLs), CD40 signaling cannot be activated by soluble CD40L but is instead activated by membrane-bound CD40L.^26^ Due to the migratory nature of both B and T cells during their encounters and the subsequent formation of T-B immunological synapses where relative sliding between two cell membranes occurs against the resistance of cross-junctional interactions, there should be ample opportunities for engaged receptor–ligand bonds, including CD40–CD40L bonds, to experience mechanical forces.^27^ Furthermore, it has been previously observed that substrate stiffness plays a role in CD40 function, with anti-CD40 functionalized on stiffer substrates leading to enhanced B cell proliferation.^28^ Given that several immune receptors on B and T cells, including TCR and BCR, are sensitive and responsive to their mechanical environment,^27,29^ we hypothesize that CD40 functions as a mechanoreceptor to receive force-modulated signals *via* its interaction with CD40L. We further hypothesize that X-HIgM mutations suppress this interaction and dysregulate CD40 mechanosignaling, contributing to the disease. To test thess hypotheses, we deployed a battery of biophysical techniques to characterize CD40’s mechanoreceptor properties and to investigate how dysregulation of CD40–CD40L interaction and mechanotransduction by X-HIgM mutations may affect B cell signaling and function.

## Results

### Force-free 2D kinetics of CD40–CD40L interactions

So far, studies on the binding of CD40L to CD40 has been limted to measuring soluble protein ectodomain interactions in the fluid phase,^30^ *i.e*., three-dimensional (3D) binding.^31^ To better mimic the physiological situation in which the interacting proteins that are expressed as membrane-bound molecules on the surfaces of two apposing cells, we apply our mechanical-based adhesion frequency assay to measure CD40–CD40L interactions across two solid surfaces, i.e., two-dimensional (2D) binding.^32–34^ These two types of binding are so-called because the affinities of 3D and 2D binding have different dimensions, volume in 3D and surface in 2D. To do this we first produced wildtype (WT) N-terminally biotinylated CD40L ectodomain protein and five X-HIgM mutants (S132W, E142G, K143A, R207A, and A208D) (Figure 1A) using a human embryonic kidney (HEK) cell mammalian expression system, which were purified and confirmed by PAGE and immunocytometry (Figures S1A-B). Of these, K143A and A208D are found in X-HIgM patients while E142G and R207A are located one amino acid distal from the natural mutants^24,35^ (Figure 1A). While these reside on the CD40L binding site for CD40, S132W is away from the binding site (Figure 1A).^36^ We then performed the adhesion frequency assay for measuring 2D kinetics^32^ using a biomembrane force probe (BFP).^37,38^ A BFP employs a micropipette-aspirated human red blood cell (RBC) with a CD40L-bearing bead (probe) attached to its apex to act as a force sensor for detecting binding to a CD40-bearing bead (Figure 1B) or a B cell (Figure 1C) (target) aspirated by an opposing micropipette (Figure S1C).^34^ The adhesion frequency (*P*_a_ = # of binding events divided by # of contacts) was measured by repeated contact cycles for each bead pair, obtaining Mean *P*_a_ ± SEM as a function of contact time (*t*_c_) using 4-5 probe-target pairs per *t*_c_value. To examine the kinetics of the interaction and directly compare among different CD40L constructs, we converted the contact time-dependent *P*_a_values to the average number of bonds per contact, 〈*n*〉 = − ln(1 − *P*_a_), based on the Poisson distribution of bonds,^32^ normalized them by the densities of both CD40 (*m*_CD40_) and CD40L (*m*_CD40L_), and plotted the resulting normalized bond number (〈*n*〉/*m*_CD40_*m*_CD40L_) *vs t*_c_ for soluble CD40-coated beads (Figure S1D) and CD40-expressing peripherial blood mononuclear cell (PBMC)-derived B cells (Figure 1D) or a germinal center-line B cell lymphoma (GCB lymphoma) Farage B cells (Figure 1E). We then fitted such data by a kinetic model,^32^ 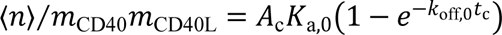, to obtain the effective 2D affinities (*A*_c_*K*_a,0_) and off-rates (*k*_off,0_) of WT CD40L and X-HIgM mutants for the CD40 ectodomain (Figures 1F-G) and for CD40 on the B cell membrane (Figures 1H and S1E). The bead-bead experiments (Figure 1B) rule out any possible confounding factors related to cellular and membrane processes involving CD40 that may be present in the bead-cell experiments (Figure 1C). Despite that the contact area *A*_c_ might vary from system to system, the consistences among WT *A*_c_*K*_a,0_ values in Figures 1F and 1H obtained from different systems indicate that the measurements on B cell surface are not confounded by such factors. Remarkably, soluble CD40 showed reduced 2D affinities for both CD40L^K143A^ and CD40L^A208D^ compared to CD40L^WT^ and reduced off-rate from CD40L^K143A^ but not from CD40L^A208D^ (Figures 1F-G). The X-HIgM mutants led to 2.2 and 2.6 (S132W), 2.6 and 1.8 (E142G), 37 and 8.2 (K143A), 17 and 7.8 (R207A) folds 2D affinity reductions for CD40 on PBMC and Farage B cells, respectively (Figure 1H), as well as few folds off-rate variations (Figure S1E) compared to CD40L^WT^.

**Figure 1.**
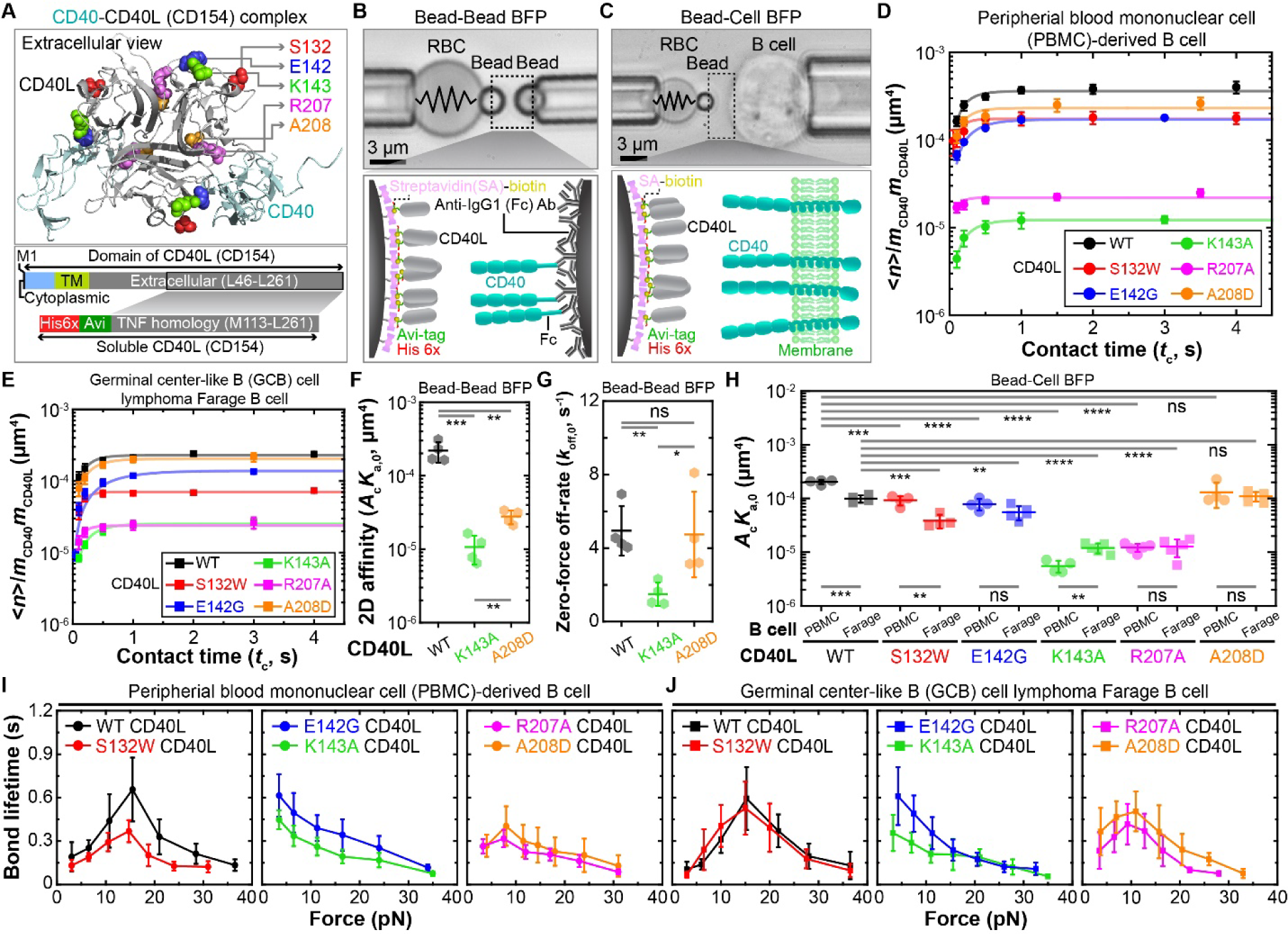
Deciphering binding parameters of CD40 interactions with CD40L constructs. (A) Crystal structure of the extracellular domains of the CD40–CD40 ligand (CD154) complex (upper, *PDB* 3QD6) and domain diagram of the soluble CD40L (lower). Colored-ball symbols highlight residues related to X-HIgM type Ⅰ (upper, indicated). The soluble CD40L used in this study was generated by truncating the full-length CD40 at residues M113-L251 with the addition of 6xHis- and Avi-tag (lower). (B and C) Photomicrographs of BFP setups for bead-bead (B) and bead-cell assays (C). A biotinylated human red blood cell (RBC) is aspirated by a micropipette and a CD40L-functionalized glass-bead (probe) is attached to its apex via biotin–streptavidin (SA) coupling (upper). The RBC-bead assembly also acts as a ultrasensitive force sensor, as indicated by the spring (left), to detect binding of CD40L with soluble CD40-Ig Fc captured by an anti-Fc antibody covalently linked to a glass bead (B) or CD40 on a B cell (C) aspirated by an opposite micropipette (right). The lower panels are composite schemes showing the interacting proteins of the two setups. Scale bar = 3 μm. (D and E) Plots of Mean ± SEM of average number of bonds per contact (⟨*n*⟩) normalized by site densities of CD40 (*m*_CD40_) and CD40L (*m*_CD40L_) *vs* contact time (*t*_c_). ⟨*n*⟩ (= − ln(1 − *P*_a_)) was converted from the adhesion frequency (*P*_a_ = # binding events divided by total # of contacts) directly measured with the indicated CD40L constructs coated on probe beads interacting with CD40-expressing PBMC (D) or Farage lymphoma (E) B cells using the BFP setup in C. Data (points, *n* = 4-5) were fitted (curves) by a previously reported model:^32^ 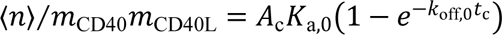 (F-H) Mean ± SD of zero-force effective 2D affinity (*A*_c_*K*_a,0_) (F and H) and off-rate (*k*_off, 0_) (G) obtained from the bead-bead (F and G) and bead-cell (H) assays. Scatters indicate individual data from each bead-bead (*n* = 4) or bead-cell (*n* = 4-5) pairs. Two-sided t-test was used to assess significance (ns > 0.05, 0.01 < ∗ < 0.05, 0.001 < ∗∗ < 0.01, 0.0001 < ∗∗∗ < 0.001, and ∗∗∗∗ < 0.0001). (I and J) Mean ± SEM of bond lifetime *vs* force plots (*n* > 35 lifetime measurements per force bin) of indicated CD40L construct-coated beads interacting with CD40-expressing PBMC (I) or Farage lymphoma (J) B cells using the BFP setup in C and the force-clamp assay shown in Figure S1C. Different panels are categorized by the different types of bond profiles: catch-slip bond for WT and S132W (lef), slip-only bond for E142G and K143A (middle), suppressed catch-slip bond for R207A and A208D (right).

### CD40 forms catch bonds with CD40L that are suppressed or eliminated by X-HIgM mutations

The effective 2D affinities and off-rates shown in Figures 1F-H and S1E were measured under the force-free condition by nature of the adhesion frequency assay,^32^ which is indicated by the subscript 0. We next asked whether, and if so, how force impacts the dissociation kinetics of CD40–CD40L bonds. To address this question, we used BFP to load single CD40–CD40L bonds with precisely controlled forces and measured the bond lifetimes under a range of constant forces (Figure S1C). Interestingly, CD40L^WT^ formed a catch-slip bond with CD40, where lifetime increases with increasing force up to ∼15 pN (optimal force) and then decreases with further increase in force; this held true for both PBMC and GCB lymphoma Farage B cells (Figures 1I-J, both with WT CD40L), suggesting that the dissociation of CD40–CD40L bond is modulated by force. This is intriguing because catch bonds have also been observed in several immunoreceptors, for instance, the TCR forms catch bonds with agonist pMHCs.^39–41^ Intuitively, catch bonds enable force to prolong engagement time and stabilize interacting molecular complexes, which allows sufficient time for downstream biological processes to proceed as needed, resulting in more protein docking and conformational changes, more enzymatic modifications, formation of more molecular assemblies (*e.g.*, signalosomes, biomolecular condensates, focal adhesions, etc.), and possibly others.

Remarkably, X-HIgM mutations to the CD40L binding site (Figure 1A) either greatly suppressed the extent of catch bonds (Figures 1I-J, R207A and A208D) or completely abrogated them, turning the CD40–CD40L interaction into slip-only bonds (Figures 1I-J, E142G and K143A); again, these results held true for both PBMC and Farage B cells, suggesting that X-HIgM mutations adversely impact the force modulation of CD40–CD40L bond dissociation, which may translate into dysregulation of CD40 signaling and downstream biological processes. The CD40L^S132W^ mutation distal to the binding site (Figure 1A) lowered the extent of catch-bond in PBMC B cells (Figure 1I, S132W) but not in Farage B cells (Figure 1J, S132W). Since for exponentially distributed lifetimes expected for single-bond dissociation, average bond lifetime equals reciprocal off-rate,^42^ we plotted the reciprocal zero-force off-rates measured from the adhesion frequency experiment *vs* the bond lifetimes measured at the lowest force bin by the force-clamp experiment, finding excellent correlation (Figures 1I-J and S1E-F), thus further confirming the short lifetimes of the catch-slip bonds (WT, S132W, R207A, and A208D) and the long lifetimes of the slip-only bonds (E142G and K143A) at low forces. It is informative that this change in bond type with the CD40L constructs tested was observed not only for membrane CD40 on the B cell surface, but also for soluble CD40 ectodomain (Figure S1G), implying that the CD40–CD40L catch bond is governed by ectodomain interactions between CD40 and CD40L.

### CD40 experiences B cell-generated endogenous forces upon engaging immobilized CD40L, which are suppressed by X-HIgM mutations

While the CD40–CD40L catch bond was demonstrated by loading the bond by a range of external forces, B cells have been observed to exert endogenous forces on BCR.^29,43,44^ We, therefore, asked whether CD40 experiences B cell-generated forces upon engagement with surface-bound CD40L in the range where the catch bond was observed. To answer this question, we employed DNA-based molecular tension probes (MTPs).^45^ An MTP consists of a DNA hairpin that unfolds at a desired tension designed by adjusting the length and guanine-cytosine content of the DNA sequence and is flanked by a fluorophore-quencher pair. One end of the hairpin is functionalized with a CD40L, anti-CD40 (positive control), or BSA (negative control) molecule and the other end is linked to a gold nanoparticle-coated surface (Figure 2A). If a B cell placed on the MTP surface exerts tension on a CD40–CD40L bond (*F*_Cell_) that is greater than the DNA hairpin force threshold (*F*_1/2_), the hairpin would (have a 50% chance to) unfold, separating the quencher (BHQ2) from the fluorophore (Atto647), thus generating a fluorescent signal to report a force of *F*_Cell_ > *F*_1/2_ on the bond (Figure 2A). Adding a single-stranded DNA (locker) complementary to the opened hairpin to prevent it from closing after bond dissociation or force removal allows the fluorescent signal to accumulate.^46^ Using this technique, we observed that both PBMC (Figure 2B) and Farage (Figure 2C) B cells efficiently spread on WT CD40L-functionalized MTP surfaces with *F*_1/2_ of 4.7, 12, and 19 pN and exerted endogenous tensions *F*_Cell_ above 4.7 pN but below 19 pN (Figures 2B-G and S2A-J). By comparison, when B cells were placed on X-HIgM CD40L mutant-tagged MTP surfaces, their spreading area and cumulative tension signals were significantly lower at all *F*_1/2_ levels regardless of the cells tested (Figures 2B-G and S2A-J). On MTP surfaces of *F*_1/2_ = 4.7 pN, PBMC and Farage B cells generated 7.2- and 11.1-fold less tension on CD40L^K143A^ and 8.8- and 19.5-fold less tension on CD40L^A208D^, respectively. To directly compare the tension signals measured using MTPs of different *F*_1/2_ values tagged by different CD40 binding molecules, we normalized all tension signals by that measured using MTP of *F*_1/2_= 4.7 pN tagged by CD40L^WT^ (Figures 2H-I). For PBMC and Farage B cells on CD40L^WT^ MTPs, the >12 pN tension signals decreased by 19.5% and 25%, respectively, whereas the >19 pN tension signals dropped by 74% and 68%, respectively, compared to the >4.7 pN tension signal (Figure 2H-I). Thus, B cells exert endogenous forces *F*_Cell_between 12 and 19 pN on CD40. This suggests a match of the active tension to the force where the lifetime of the CD40–CD40L^WT^ bond peaks as it transitions from the catch regime to the slip regime (i.e., optimal force). By comparison, the tension signals of our positive control (anti-CD40) decreased precipitately with the MTP *F*_1/2_ from 4.7 to 12 and 19 pN (Figures 2H-I), which correlates with the slip-only behavior of the CD40–anti-CD40 bond (Figures S2K-L). These data are remarkable, as they suggest a potential mechanobiological mechanism in which CD40 and/or CD40L may be evolved to enable the formation of a catch-slip bond with the force where the bond lifetime peaks to match the endogenous force generated by the cell’s cytoskeleton and motor machinery.

**Figure 2.**
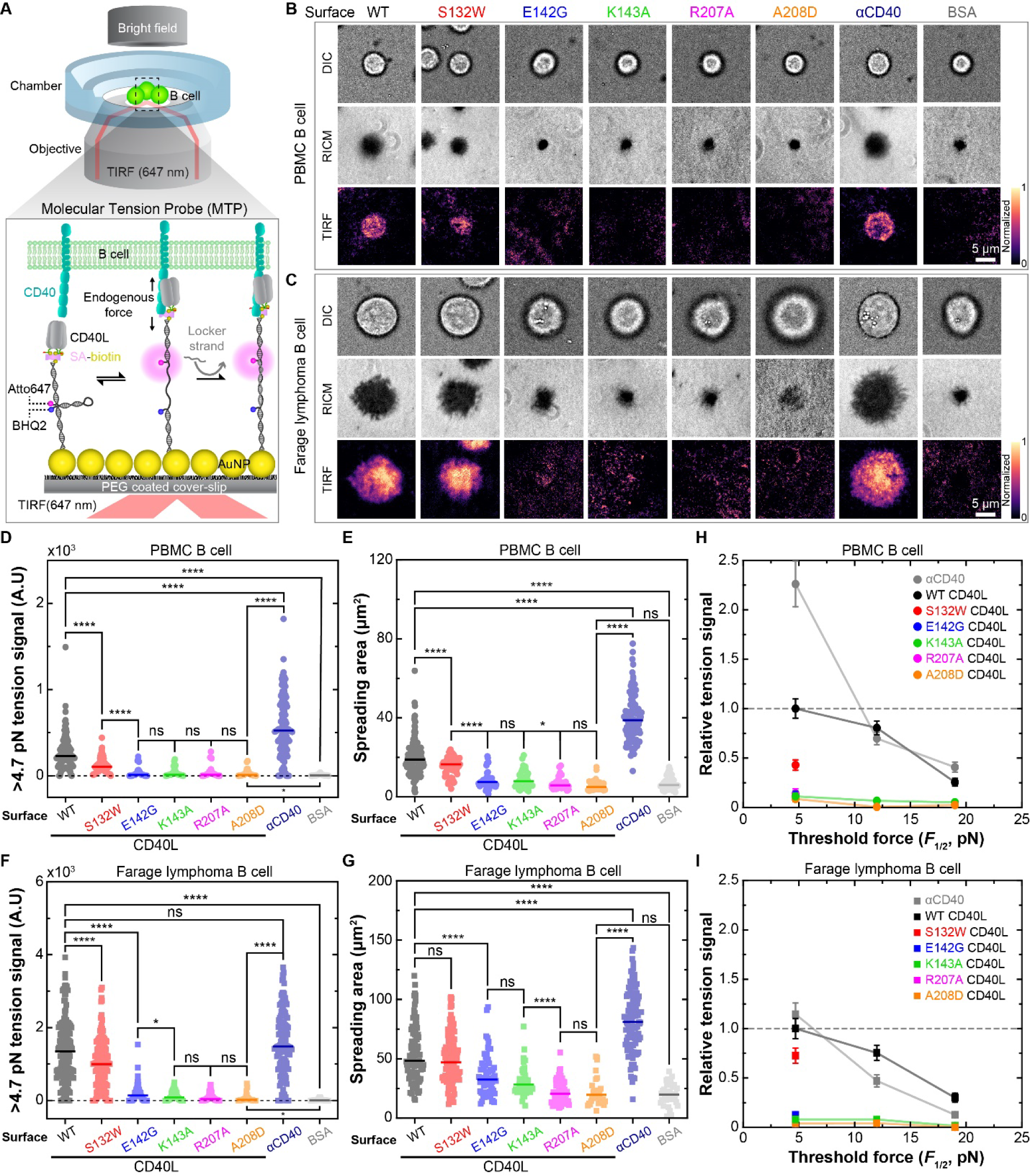
CD40 experiences B cell-generated forces upon engaging immobilized CD40L, which are suppressed by X-HIgM mutations. (A) Schematics of molecular tension probe (MTP) setup and its working principle.^45^ An MTP consist of a DNA hairpin whose two closed ends are conjugated with a fluorophore (Atto647) and quencher (BHQ2) pair. One end is linked to a CD40L (or an anti-CD40 antibody for positive control or BSA for negative control) and the other end is linked to a gold nano paticle (to further quench the fluoreophore) bound to a polyethylene glycol (PEG)-coated glass surface (lower left). The hairpin unfolds at a given tension depending on the length and %GC of the DNA sequence. Placed on the stage of an inverted microscope, CD40L on the MTP surface interacts with CD40 on B cells (upper), which exert tension on the CD40–CD40L bonds (*F*_Cell_). If *F*_Cell_ is greater than the DNA hairpin force threshold (*F*_1/2_), the hairpin would unfold, separating BHQ2 from Atto647, thus generating a fluorescent signal (lower middle). Adding a ssDNA (locker) complementary to the opened hairpin to the system prevents it from closing after bond dissociation and force removal, which allows the fluorescent signal to accumulate^46^ (lower right). (B and C) The representative images of differential interference contrast (DIC, 1^st^ row), reflection interference contrast microscopy (RICM, 2^nd^ row), and total internal reflection fluorescence normalized tension signal (TIRF, 3^rd^ row) from CD40 of PBMC (B) and Farage B cell (C) interacting with indicated CD40L constructs, anti-CD40 antibody, and BSA tagged MTP surface. Scale bar = 5 μm. (D-G) Mean and individual data points of >4.7pN tension signals (D and F) and spreading areas (E and G) of PBMC (D and E) and Farage (F and G) B cells (*n* = 25-200) on MTP surfaces functionalized indicated CD40L constructs, anti-CD40 antibody, and BSA. Two-sided t-test was used to assess the significance of differences between groups (ns > 0.05, 0.01 < ∗ < 0.05, 0.001 < ∗∗ < 0.01, 0.0001 < ∗∗∗ < 0.001, and ∗∗∗∗ < 0.0001). (H and I) Mean ± SEM of relaive tension signals obtained from placing PBMC (H) and Farage (I) B cells on the indicated MTP surfaces. The relative tensions are normalized by the >4.7 pN tension MTP surface conjugated with WT CD40L (*n* = 25-262 cells).

### Endogenous force supports B cell spreading on CD40L and limiting force suppresses CD40 signaling

Given our finding that B cells generated forces on CD40–CD40L bonds that match their force-dependent dissociation kinetics, we asked whether such endogenous forces are important to CD40-mediated B cell signaling and function. To address this question, we capped the force magnitude that a B cell could exert on any single CD40 molecule and assessed the ability of B cells to spread on CD40L and signal. This was done using the tension gauge tether (TGT) technology.^47^ TGT is similar to MTP in that both take advantage of the property of double-stranded (ds) DNA oligos to dissociate above a tunable force threshold (Figure 3A). However, MTP is a single strand (ss) DNA that folds into a hairpin structure, unfolds by an above threshold force, and refolds reversibly after force removal unless another complementary ssDNA hybridizes with the unfolded MTP to lock it in the open configuration (Figure 2B). By comparison, TGT uses dsDNA with open ends, hence will rupture irreversibly by above threshold forces (Figure 3A).

**Figure 3.**
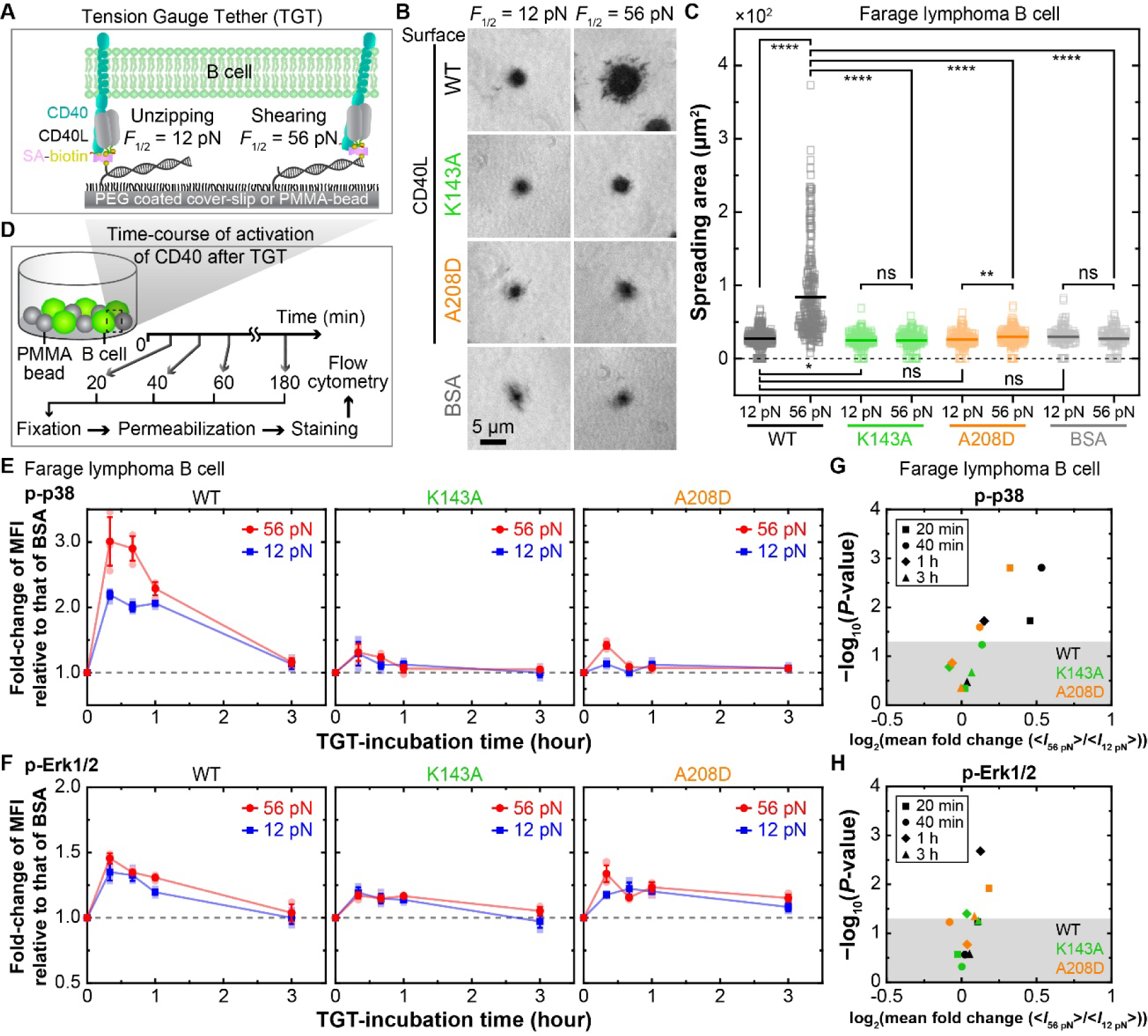
Endogenous forces support B cell spreading on CD40L and limiting force suppresses CD40 signaling. (A) Schematic of tension gauge tether (TGT) setup and its working principle. TGTs consist of open-ended dsDNAs with one strand linked to a PEG-coated glass surface *via* azide-DBCO conjugation and the other strand tagged to a CD40L molecule *via* SA-biotin coupling. If endogeneous forces exerted on CD40–CD40L bonds exceed the designed threshold, the dsDNAs dissociate, limiting the amount of forces that the B cell can experience through CD40 even if CD40L remains engaged. TGT’s force threshold depends not only on the length and %GC of the DNA sequence but also on the mode of dissociation. Using the two strands from the same end to link the dsDNA to the surface and CD40L forces dissociation to follow a “unzipping” mode, giving rise to a *F*_1/2_ = 12 pN (left), whereas lining the surface and CD40L via different ends of the dsDNA necessitates dissociation to follow a “shearing mode”, yielding a *F*_1/2_= 56 pN (right). (B) Representative RICM images of Farage cell spreading on TGT of 12 pN (left) or 56 pN (right) force threshold tagged with indicated CD40L constructs as well as BSA (negative control). Scale bar = 5 μm. (C) Mean with individual data points (*n* = 116-309) of area of Farage cell spreading on TGT surfaces functionalized with indicated molecules. Two-sided t-test was used to assess significance of differences between groups (ns > 0.05, 0.01 < ∗ < 0.05, 0.001 < ∗∗ < 0.01, 0.0001 < ∗∗∗ < 0.001, and ∗∗∗∗ < 0.0001). (D) Experiment scheme for measuring intracellular signaling after stimulation by TGT beads. Azide-PMMA (poly(methacrylic acid)) beads are functionalized TGT (*F*_1/2_ = 12 or 56 pN) tagged with CD40L or BSA *via* SA-biotin coupling. After co-culturing with TGT beads for various incubation times, B cells were fixed, permeablized, stained with antibodies against phorsphorylated signaling proteins, and analyzed by flow-cytometry. (E and F) Mean ± SD with individual data points (*N* = 3 with n > 10,000 cells) of fold-change of mean fluorescence intensity of PE (MFI) normalized by that of BSA for phopho-p-38 (p-p38) (E) and phospho-Erk1/2 (p-Erk1/2) (F) in Farage cells stimulated with TGT-tagged CD40L constructs (indicated on top of each panel) are plotted *vs* incubation time. The capacities to stimulate B cell signaling of two types of TGT beads are compared, with respective *F*_1/2_= 12 pN (blue) and 56 pN (red). (G and H) Log plots of relative change of *P*-value *vs* mean fold change of MFI of p-p38 (G) and p-Erk1/2 (H) between Farage cells stimulated by TGT with 12 pN and 56 pN force thresholds. Different points represent different TGT-incubation times and different colors indicate different CD40L constructs. Shaded area represents region of *P*-value > 0.05 above which indicates region of statistical significance.

Given our measured bond lifetime *vs* force curves (Figures 1I-J), we chose two TGT designs that detach at threshold forces of 12 and 56 pN, determined by their unzipping and shearing rupture modes, respectively (Figure 3A).^47^ When Farage cells were placed on 56-pN TGT surfaces functionalized with CD40L^WT^, allowing up to 56 pN endogenous forces to exert on CD40–CD40L bonds, the cells spread nicely (Figures 3B-C). In contrast, when the TGT threshold force was lowered to prevent >12 pN force on CD40–CD40L bonds, cell spreading was greatly reduced, indicating that force is important for B cell spreading on CD40L. Compared to WT, X-HIgM mutations to CD40L failed to support appreciable spreading of Farage cells, showing statistically indistinguishable spreading areas on CD40L^K143A^-functionalized 12- and 56-pN TGTs but significantly smaller spreading areas on CD40L^A208D^-functionalized 12-than 56-pN TGT (Figures 3B-C).

B cells spread on FDCs to capture membrane-bound antigens^48^ and on T_FH_ cells to form immunological synapse,^49^ both of which are important.^50^ Cell spreading has an active component that involves signaling.^51–53^ As a first step towards identifying specific molecules downstream of CD40 signaling that might require B cell endogenous forces on CD40, we developed a bead-based TGT assay coupled with flow cytometry (Figure 3D) to measure phosphorylation of p38 and Erk1/2, two kinases known to be important parts of CD40-induced signaling pathways.^35,54–56^ B cells were co-incubated with 12-μm-diameter beads functionalized with CD40L-tagged 12 or 56 pN TGTs, and subsequently harvested at different timepoints to measure phosphorylation of p38 and Erk1/2. Of note, the size of beads functionalized with TGT should be similar or even larger than the cell size in order to instigate a contact area for maximal interactions. The phosphorylation of both intracellular molecules showed a biphasic timecourse that peaked at ∼20 min, with levels at all timepoints higher when Farage cells were allowed to exert up to 56 pN forces on CD40–CD40L bonds than when the forces were capped to <12 pN (Figures 3E-F and S3A). The differences in p38 phosphorylation were significant at 20, 40, and 60 min (Figures 3E and 3G). Erk1/2 phosphorylation showed less pronounced changes and was significantly reduced by <12-pN relative to <56-pN TGT at 60-min timepoint only (Figures 3F, H and S3A). Thus, force differentially impacts p38 and Erk1/2 signaling, with the p38 pathway more dependent on force than the Erk1/2 pathway. Trends equivalent to those described above for Farage B cells were observed for PBMC B cells (Figures S3B-F), indicating that these results are not artifacts of lymphoma cells.

Consistent with their suppressed force-dependent kinetics (Figures 1I-J) and their inability to exert higher forces (Figures 2B-G and S2A-J), phosphorylation of p38 and Erk1/2 for cells incubated with TGTs tagged with X-HIgM CD40L mutants was greatly reduced, with CD40L^K143A^ (which forms a slip-only bond with CD40) being reduced more than CD40L^A208D^ (which forms a suppressed catch-slip bond with CD40) (Figures 3E-H and S3). Also, no significant differences in phosphorylation timecourses of p38 and Erk1/2 were detected between B cells whose forces on CD40–CD40L^K143A^ bonds were limited to be <12 pN and those that allowed up to 56-pN forces (Figures 3E-H and S3). By comparison, we observed higher phosphorylation of these two signaling molecules when B cells were allowed to exert up to 56 pN forces on CD40–CD40L^A208D^ bonds than when the endogenous forces were limited to be <12 pN (Figures 3E-H and S3A-D). Taken together, these results indicate that endogenous forces on CD40–CD40L bonds are necessary for B cell spreading and differentially enhance CD40 proximal signaling. X-HIgM mutations abolish spreading and diminish signaling due to their inability to form long-lived bonds at higher forces.

### External force on CD40–CD40L bonds amplifies CD40 signaling

Our finding that limiting endogenous forces on CD40 suppresses B cell spreading and signaling prompted us to ask whether applying an external force to CD40 would enhance B cell signaling and function. After all, B cells exerts endogeneous force on CD40 and a reaction force of equal magnitude but opposite direction would be acted back to B cells through CD40 by virtue of Newton’s third law. To answer this question in a high throughput fashion, we developed a parallel magnetic force activation (PMFA) assay (Figures 4A and S4A-D), modified from a published system.^57^ Our multi-well plate-based assay takes advantage of a 3D-printed plate lid (Figure S4A) that can house two magnets placed in the antiparallel configuration above each well, which can apply magnetic force on paramagnetic beads bearing CD40L bound to CD40 on B cells underneath the magnets (Figure S4B). While accounting for the size constraint of cell media in each well, each magnet pair is placed as low as possible inside the wells, allowing 20-pN average force per bead (〈*F*_M_〉) and a peak force of ∼25pN at the center of the well (Figure S4C). To constrain the bead-bounded cells to the well center where the desired range of magnetic forces are focused, we seeded cells and beads on a poly-L-lysine (PLL)-coated 5-mm diameter coverslip placed in the center of each well (Figure 4A and S4B). Cells seeded on the coverslip naturally concentrated towards its center, with a distribution closely matching the distribution of the magnetic force exerted on beads (Figures S4D), thus allowing force to be evenly distributed on as many cells as possible. Assuming the applied force on each bead to be shared evenly by all CD40–CD40L bonds present, we adjusted the CD40L site density coated on beads to achieve the desired steady-state average number of bonds (〈*n*〉) per bead, based on the CD40 expression on B cells and the 2D affinity measured by BFP (Figure 1H), to control the force per bond (*f*_M_ = 〈*F*_M_〉/〈*n*〉) to match the endogenous force range (∼ 10-20pN) (Figures S4E-F).

**Figure 4.**
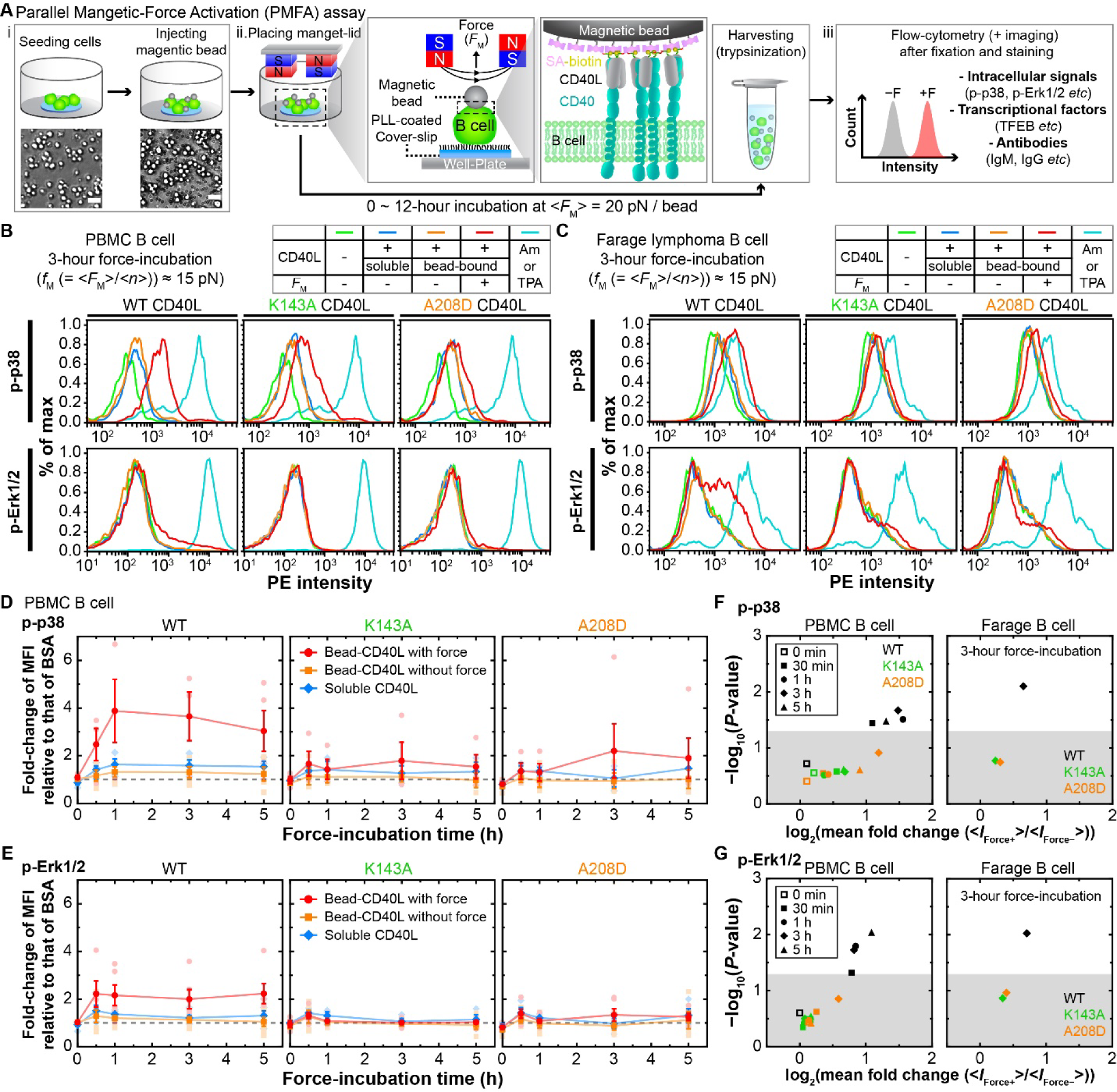
Exogenous forces on CD40–CD40L bonds amplify CD40 signaling. (A) Working principle and procedure of the parallel magnetic force activation (PMFA) assay. i. Schematic (top) and representative images (bottom) of B cells placing on PLL-coated circular coverslip (5-mm in diameter) at the well center (left) onto which magnetic beads (2.8-μm in diameter) funcionalized with CD40L constructs are added (right). Scale bar = 40 μm. ii. Zoom-out (far left) and zoom-in (2^nd^ left) views showing how an average force of 〈*F*_M_〉 ∼ 20 pN/bead is applied on the bead by placing a magnet-lid on the 24-well plate, with an anti-parallel pair of 5-mm cubic neodymium magnets added to each well. This force is supported by the CD40–CD40L bonds (2^nd^ right). After 0-12 hrs of incubation, cells are harvested (far right). iii. Then, cells are fixed, permeabilized, and stained with fluorescence-conjugated antibodies against phosphorylated intracullar signaling molecules, transcriptional factor and nucleus, or membrane-bound Ig types. Cells are then analyzed by flow-cytometry. (B and C) Keys (tables) and representative flow-cytometry data (plots of normalized frequency of cells *vs* PE-intensity) of p-p38 (top row) and p-Erk1/2 (bottom row) of PBMC (B) and Farage (C) B cells stimulated by soluble or bead-bound CD40L constructs (indicated on top of each panel) without or with 3-h force application. Also shown are data of negative (no stimulation) and positive (AM or TAP stimulation) controls. (D and E) Mean ± SEM with individual data points (*N* ≥ 3 with n > 10,000 cells) of fold-change of MFI relative to that of BSA for p-p-38 (D) and p-Erk1/2 (E) in PBMC B cells stimulated by soluble (blue) or bead-coated CD40L constructs (indicated on top of each panel) with (red) or without (orange) force are plotted *vs* incubation time. (F and G) Log plots of relative change of *P*-value *vs* mean fold change of MFI of p-p38 (F) and p-Erk1/2 (G) in PBMC (left) and Farage (right) B cells between stimulations by bead-bound CD40L without and with force. Different scatter points in each plot represent different force-application times and different colors indicate different CD40L constructs. Shaded area represents *P*-value > 0.05 above which indicates region of statistical significance.

Enabled by the PMFA technological platform, we stimulated large numbers of B cells in parallel in the presence or absence of exogenous force on CD40–CD40L bonds, harvested them at various timepoints, stained them with fluorescence antibodies against phosphorylated p38 and Erk1/2, and analyzed them by flow cytometry (Figure 4A). Phosphorylation of p38 and Erk1/2 in either PBMC or Farage B cells stimulated by soluble and by bead-coated CD40L in plates without magnets on their lids were statistically indistinguishable, which were barely higher than those in unstimulated cells (negative control) but much lower than those in anysomicin (AM) or tetradecanoylphorbol acetate (TPA)-stimulated cells (positive control) for p38 and Erk1/2, respectively (Figures 4B-C). Mounting magnets on the plate lids greatly right-shifted the fluorescence histograms, indicating that applied force on CD40 upregulated p38 and Erk1/2 phosphorylation in both PBMC or Farage B cells (Figures 4B-C). Compared to removal of the 12-pN cap on endogenous forces by TGT (Figures 3E-F), applying exogenous forces accelerated and sustained the phosphorylation kinetics of p38 and Erk1/2, which peaked at 1 h post-stimulation and were maintained up to at least 4 more hours (Figures 4D-G).

Again, the X-HIgM CD40L mutants showed defects in force-induced p38 and Erk1/2 phosphorylation, with greater deficiencies for CD40L^K143A^ than CD40L^A208D^, indicating eliminating catch bonds is more defective than suppressing catch bonds (Figures 4B-E). The force-dependent enhancement of signaling for CD40L^WT^ and its reduction by X-HIgM CD40L mutants could be revealed by comparing the mean phosphorylation fold-change of cells with and without force to the significance level (*P*-value) at each time point (Figures 4F-G). Similarly, trends equivalent to those described above for PBMC B cells were observed for Farage cells (Figure S4G), indicating that these results are not affected by B cell types. Moreover, as expected, all soluble CD40L constructs tested showed statistically indistinguishable results compared to those tested under no-force conditions (Figures S4H-I). Only CD40L^WT^ showed significant force-enhancement as all but the 0 timepoint WT data appear in the p < 0.05 region (white), whereas data for both X-HIgM mutants appear in the p > 0.05 region (gray), consistent with their shorter bond lifetimes (Figures 1I-J), less endogenous force on CD40 (Figures 2D-G), and inability to support spreading and signaling even when endogenous forces were not limited to <12 pN (Figures 3E-H). Taken together, these observations demonstrate how the force on CD40–CD40L bonds can enhance CD40 signaling and how X-HIgM mutations impair this force-induced signaling enhancement, regardless of B cell types and the source of the force.

### Force on CD40–CD40L bonds amplifies BCR-dependent translocation of TFEB

BCR ligation in the absence of co-stimulation by T cells induces B cell death by apoptosis but the underlying mechanism remained mysterious.^2^ Recently, the BCR-induced nuclear translocation of the cytosolic transcription factor EB (TFEB) was identified as a key process that governs apoptosis of B cells.^58^ Nuclear TFEB drives B cell apoptosis through the expression of pro-apototic Bcl-2 family members.^58^ At the same time, TFEB is involved in rescuing B cells from apoptosis *via* the transcriptional upregulation of receptors that are involved in receiving secondary salvage signals (*e.g*., cytokine receptors, homing receptors, MHC-II). CD40 ligation interferes with these signal circuits in two ways. One the one hand, CD40 activation antagonizes the TFEB-induced expression of pro-apoptotic Bcl2 family members by triggering the expression of the anti-apoptotic family member Bcl-xL, which consequently halts the apoptotic process. On the other hand, CD40 signaling sustains the nuclear residency of TFEB, thereby further promoting the expression of TFEB-controlled salvage elements and concomitant B cell survival.^58^

Since force on CD40 affected bond kinetics and signaling, we hypothesized that force on CD40-CD40L bonds would promote CD40-induced TFEB translocation. To test this hypothesis, we combined our PMFA assay with imaging flow cytometry to measure the localization of TFEB following BCR stimulation with soluble anti-IgM and CD40 stimulation with magnetic bead-bound CD40L in the presence or absence of force using a similarity score to quantify the overlap between the TFEB (FITC) and the nucleus (7-AAD) staining (Figures 5B). Due to the lack of BCR expression on Farage cells, we switched to Ramos B cells after confirming that their force-dependent interaction kinetics of CD40 with CD40L and its X-HIgM mutants were similar to Farage and PBMC B cells (Figures S5A-C). Consistent with a previous study,^58^ Ramos cells stimulated with soluble anti-IgM had markedly increased translocation of TFEB with a higher portion of cells with nuclear TFEB as indicated by the higher similarity score, compared to cells without BCR stimulation whose TFEB was mostly localized in the cytoplasm with a low similarity score (Figures 5B-E and S5D-F). We next compared the effect of co-stimulation by soluble or bead-bound CD40L in the absence and presence of exogenous force on the BCR-stimulated TFEB translocation by normalizing the similarity score and proportion of cells with nuclear TFEB to that of B cells stimulated with soluble anti-IgM alone (Figures 5D-E). Remarkably, while both soluble and bead-bound CD40L showed statistically indistinguishable TFEB translocation in the absence of force, applying force to CD40–CD40L bonds significantly increased the similarity score and the portion of cells with nuclear TFEB (Figures 5D-E). In sharp contrast, X-HIgM CD40L mutants showed no force-induced increase in TFEB translocation when compared to BCR stimulation alone. These findings indicate that force on CD40 is required for the additionial enhancement of TFEB translocation co-stimulated by soluble anti-IgM and bead-coated CD40L (relative to soluble anti-IgM stimulation alone), and that X-HIgM CD40L mutants do not support such additional enhancement.

**Figure 5.**
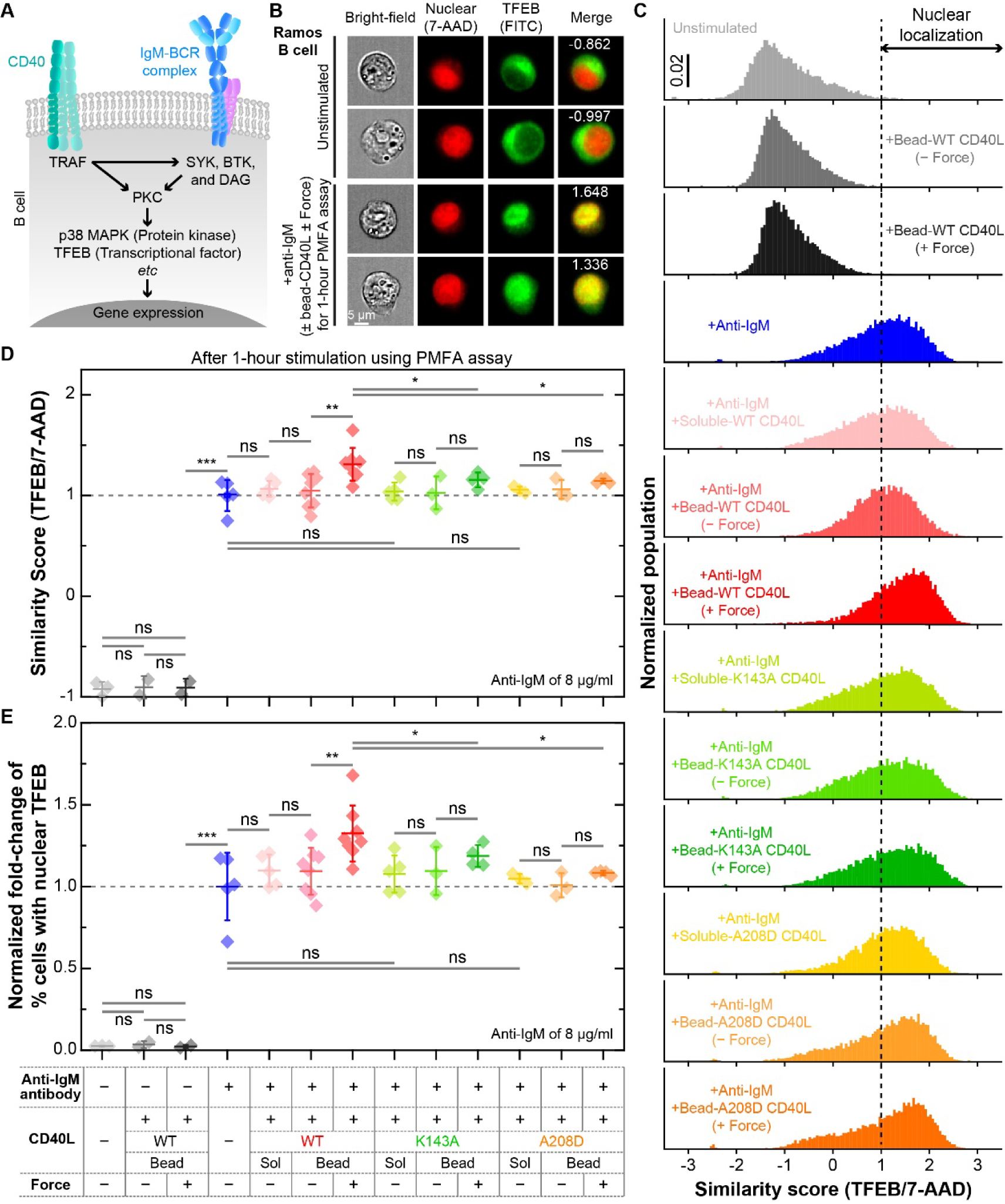
Force on CD40 –CD40L bonds amplifies BCR-induced translocation of TFEB. (A) Brief summary of B cell signaling downstream of CD40 and BCR. (B) Representative bright-field (1^st^ column) and fluorescence (2^nd^-4^th^ columns) images of image flow cytometer without (first two rows) and with (last two rows) PMFA-exerted force on CD40-CD40L bonds using Ramos B cells. Nucleus and transcriptional factor EB (TFEB) are stained by 7-AAD (2^nd^ column) and FITC (3^rd^ column), respectively. The yellow color in the merged image (4^th^ column), resulting from overlapping of the red and green fluorescences, reveals the translocation of TFEB into the nucleus, as quantified by the similarity scores (numbers in the merged images). (C) The normalized populations of similarity scores for different conditions as indicated by different colors in the different panels (in descending order): unstimulated (light gray), bead-coated CD40L^WT^ without force (dark gray), soluble anti-IgM antibody (blue), soluble anti-IgM plus soluble CD40L^WT^ (pink), soluble anti-IgM plus bead-coated CD40L^WT^ without force (light red), soluble anti-IgM plus bead-coated CD40L^WT^ with force (dark red), soluble anti-IgM plus soluble CD40L^K143A^ (light green), soluble anti-IgM plus bead-coated CD40L^K143A^ without force (green), soluble anti-IgM plus bead-coated CD40L^K143A^ with force (dark green), soluble anti-IgM plus soluble CD40L^A208D^ (yellow), soluble anti-IgM plus bead-coated CD40L^A208D^ without force (light orange), and soluble anti-IgM plus bead-coated CD40L^A208D^ with force (orange). (D and E) Mean ± SD with individual data points (*N* ≥ 3 with n > 10,000 cells) of fold-change of similarity score (D) and % cells with nuclear TFEB normalized by anti-IgM only data (E). Different colors indicate different conditions (matched with those in C). Two-sided t-test was used to assess statistical significance in differences among conditions (ns > 0.05, 0.01 < ∗ < 0.05, 0.001 < ∗∗ < 0.01, and 0.0001 < ∗∗∗ < 0.001).

### Force on CD40–CD40L bonds promotes antibody class-switch and allows discrimination between WT CD40L and X-HIgM mutants

To put our hypothesis that mechanotransduction governs CD40 function and dys-mechanoregulation underlies X-HIgM to a final test, we examined whether, and if so, how force affects long-term B cell function during GC formation, B cell differentiation, and antibody class switch. This is especially important in light of the observed defects in X-HIgM CD40L mutants’ ability to support CD40 mechanotransduction, which correlates with their known defects in Ig class switch recombination (CSR) leading to X-HIgM pathology. To study the effect of force on such B cell functions, we mimicked the GC reaction by an *in vitro* assay that allows us to induce CD40L-dependent B cell activation, differentiation, and CSR by co-culturing PBMC B cells with CD40L-coated beads in the presence of IL-4, IL-21 and BAFF,^59–61^ and combining this with or without force application on CD40–CD40L bonds by the PMFA technique (Figures 6A and S6A-B). After force application for the initial 6 h followed by prolonged co-culture for 8 days, B cell phenotypes and Ig types were measured by flow cytometry (Figures S6A-B) with appropriate fluorescence minus one (FMO) controls.^59^ Using this setup, no significant changes in the total CD19^+^ population, CD19^+^CD27^+^CD38^+^ activated B cell-like fraction, or CD19^+^CD27^+^CD38^−^ memory-like B cell (MBC) fraction of B cells were observed when comparing WT CD40L to X-HIgM CD40L mutants in the presence or absence of force (Figures 6B-C and S6C-D). These markers were used based on recent reports of *in vitro* human B cell cultures.^59^ For the entire CD19^+^ B cell populations, the IgM^+^ portions were similar across all CD40L constructs with or without force (Figures 6D and S6E, IgM^+^ in entire CD19^+^). By comparison, the IgG^+^ portion was significantly higher for CD40L^WT^, and significantly lower for X-HIgM CD40L mutants, in the presence compared to in the absence of force, despite that without force CD40L induced lower IgG^+^ portion than its X-HIgM mutants (Figures 6D and S6E, IgG^+^ in entire CD19^+^).

**Figure 6.**
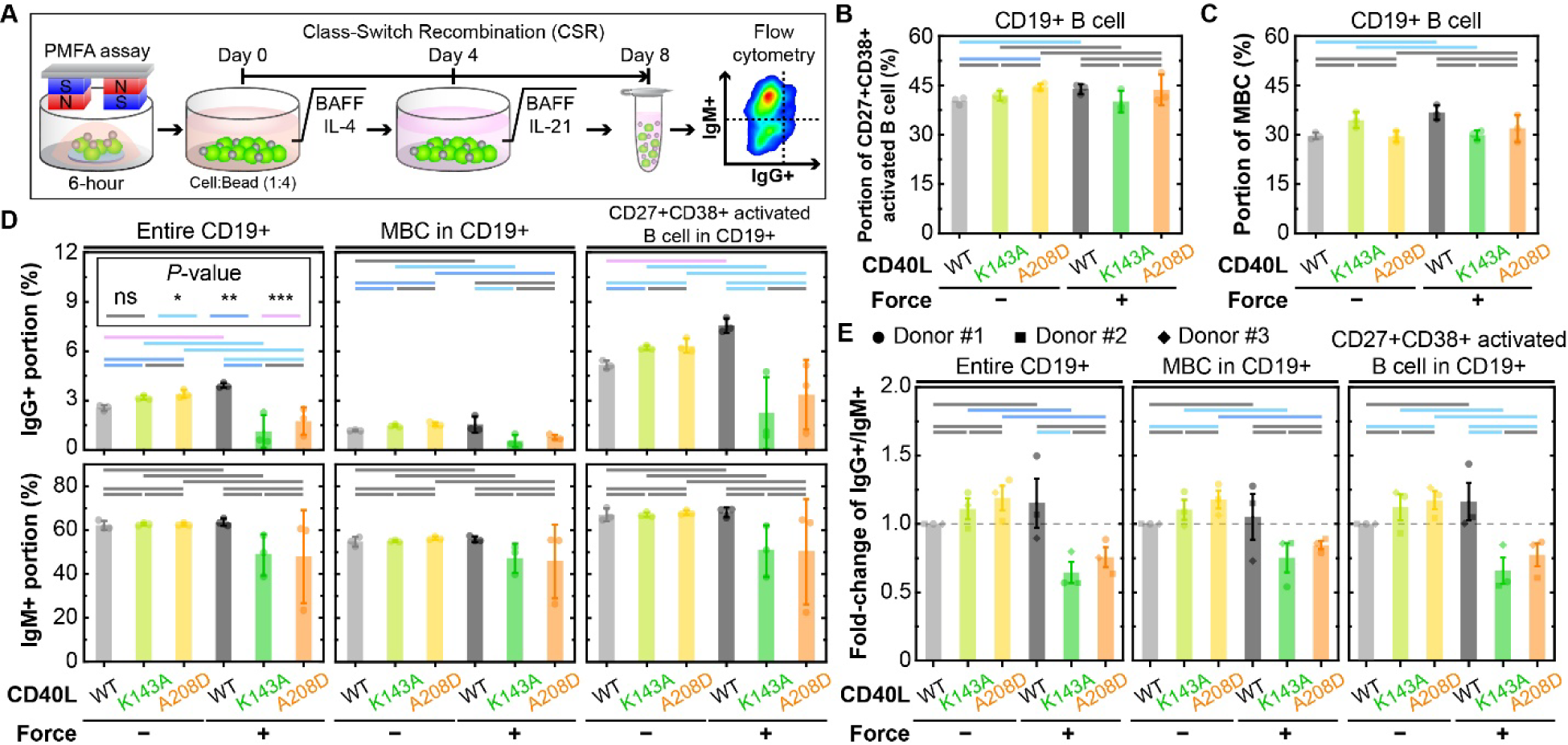
Force on CD40–CD40L bonds promotes antibody class-switch for WT CD40L and impairs class-switch for X-HIgM mutants. (A) Schematic of procedure and working principle for measuring class-switch recombination with the PMFA assay. After 6-hour force-application, cells are incubated with cytokines for 8 days (soluble BAFF and IL-4 for the first 4 days and soluble BAFF and IL-21 for the last 4 days). At day 8, cells are harvested, fixed, permeabilized, and stained to measure antibodies expressed on the membrane by flow-cytometry. (B) Mean ± SD with individual data points (*N* = 3 with n > 10,000 cells) of CD19^+^CD27^+^CD38^+^ activated B cell subpopulation among the entire population of CD19^+^ B cells from a repsentative donor stimulated by the indicated bead-bound CD40L constructs with and without force. (C) Mean ± SD with individual data points (*N* = 3 with n > 10,000 cells) of CD19^+^ Memory B cell (MBC) subpopulation among the entire population of CD19^+^ B cells from a repsentative donor stimulated by the indicated bead-bound CD40L constructs with and without force. (D) Mean ± SD with individual data points of IgG^+^ (top row) and IgM^+^ (bottom row) portions in all CD19^+^ B cells (1^st^ column), MBC (2^nd^ column), and CD19^+^CD27^+^CD38^+^ activated B cells (3^rd^ column) from a repsentative donor stimulated by the indicated bead-bound CD40L constructs with and without force. (E) Mean ± SEM (N = 3) of force-stimulated fold-change of IgG^+^ to IgM^+^ ratio normalized by the data of WT CD40L stimulation without force in all CD19^+^ B cells (left), MBC (middle), and CD19^+^CD27^+^CD38^+^ activated B cells (right) from three donors stimulated by the indicated bead-bound CD40L constructs with and without force. Two-sided t-test was used to assess statistical significance in differences among groups (ns > 0.05, 0.01 < ∗ < 0.05, 0.001 < ∗∗ < 0.01, and 0.0001 < ∗∗∗ < 0.001).

To obtain further support to the above results, we analyzed the IgM^+^ portions in the MBC and activated B cell-like subsets, finding no differences among CD40L constructs in the absence of force (Figures 6D and S6E, IgM^+^ of MBC and CD19^+^CD27^+^CD38^+^ activated B cell), akin to the result obtained from analyzing the overall B cell population (Figure 6D, IgM^+^ in entire CD19^+^). Similar to the overall B cell population, in the presence of force the IgG^+^ portion of activated B cell-like subsets was significantly increased by CD40L^WT^ and significantly decreased by X-HIgM CD40L mutants (Figure 6D, IgG^+^ in activated B cell-like subsets). The IgG^+^ portion of MBC followed similar but less pronounced trends (Figures 6D and S6E, IgG^+^ in MBC CD19^+^). The data in Figure 6D obtained from PBMCs of a representative donor were confirmed by repeated experiments using two other donors (Figures 6E and S6E-G). Interestingly, the data not only show that force upregulates Ig class switch induced by CD40L^WT^ and downregulates Ig class switch induced by X-HIgM CD40L mutants, but also suggest that force enables CD40 to distinguish between CD40L^WT^ and its X-HIgM mutants in terms of their ability to induce the Ig class switch, recapitulating the pathology of X-HIgM. The inability of CD40 to discriminate between CD40L^WT^ and X-HIgM CD40L mutants in the absence of force demonstrates that force on CD40 leads to enhanced Ig class switch and is required to reproduce the detrimental effects of the CD40L mutations on Ig class switch as observed in X-HIgM patients, shedding lights on the disease mechanism.

### Effective 2D affinity or bond lifetime at optimal force predicts B cell signaling outcome and functions

After obtaining six sets of data using four mechanobiological techniques, we asked the question of whether there are, and if so, what are the underlying causative relationships among these data. Whereas the connection from B cell signaling (*e.g.*, phosphorylation of p38 and Erk1/2 as well as TFEB translocation) to function (*e.g.*, Ig class-switch) may be intuitively conceivable, their link to CD40–CD40L binding parameters seems less appearent. Our work first extended the binding parameters from the 3D ones measured in the fluid phase to their 2D counterparts measured *in situ* at the B cell membrane, including the effective 2D on-rate *A*_c_*k*_on_, the zero force off-rate *k*_off,0_, and their ratio, the effective 2D affinity at zero-force *A*_c_*K*_a,0_ = *A*_c_*k*_on_⁄*k*_off,0_ (Figures 1D-h and S1D,E). Since the data in Figures 2-6 and S2-S6 suggest the importance of force, we also examined the force-dependent bond lifetime 〈*t*_*b*_〉(*f*). This is a natural extension of the zero-force reciprocal off-rate 1⁄*k*_off,0_ (cf. Figure S1F) by including the force modulation of bond dissociation. To include the effective 2D on-rate *A*_c_*k*_on_ that is also affected by the X-HIgM mutations, we also extended the zero-force effective 2D affinity from a value to a curve by replacing 1⁄*k*_off,0_ by 〈*t*_*b*_〉(*f*) the affinity definition to define a “force-dependent effective 2D affinity” *A*_c_*K*_a_(*f*) = *A*_c_*k*_on_〈*t*_*b*_〉(*f*) (Figure 7A).^62^ It is evident from the *A*_c_*K*_a_ ratio *vs f* curves that force amplifies the differential affinities of CD40 between CD40L^WT^ and the X-HIgM CD40L mutants until reaching ∼15 pN, the optimal force where the lifetime of the CD40–CD40L^WT^ catch-slip bond peaks (Figure 7B). This is similar to the observation that catch bond amplifies TCR discrimination of agonist vs antagonists.^39,63^

**Figure 7.**
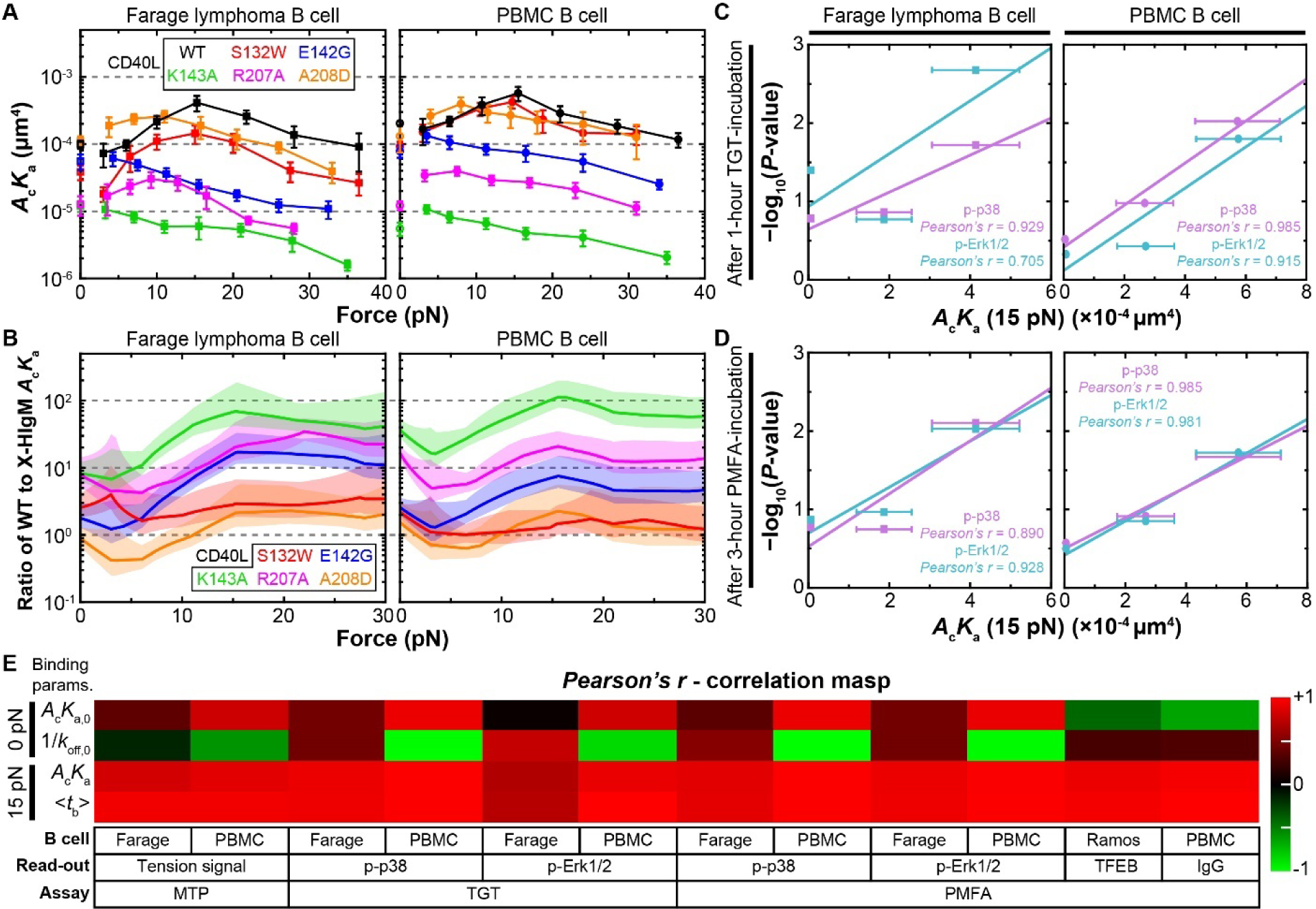
Effective 2D affinity and bond lifetime at optimal force predict B cell signaling and functions. (A) Force-dependent 2D affinities (*A*_c_*K*_a_(*f*) = *A*_c_*k*_on_〈*t*_*b*_〉(*f*), calculated using *A*_c_*k*_on_ from Figure S1E and 〈*t*_*b*_〉(*f*) from Figure 1I-J)^62^ of CD40 for the indicated CD40L constructs at the membrane of Farage (left) and PBMC (right) B cells *vs* force *f*. (B) Ratios of *A*_c_*K*_a_(*f*) of WT CD40L to those of X-HIgM CD40L mutants *vs* force *f*, showing how force amplifies the differential affinities between WT and X-HIgM CD40Ls in Farage (left) and PBMC (right) B cells. (C and D) Semi-log plots of correlation of *A*_c_*K*_a_(15 pN) of CD40 interaction with CD40L constructs with the *P*-value (from Figure 3G-H and Figure 4F-G) quantifying the statistical significance in the differences in singaling between groups (evaluated by p-p38 and pErk1/2 levels) in Farage (first column) or PBMC (last column) B cells stimulated by the same CD40L constructs under force application at two timepoints (indicated) in TGT (C) and PMFA (D) experiments. Data (points) are presented as Mean ± SEM (calculated using the Gaussian error propagation law for *A*_c_*K*_a_(15 pN), *n* = 3-4 bead-cell pairs for *A*_c_*k*_on_ and ∼35 measurements for 〈*t*_b_〉(15 pN)) and fitted by a straight line for each group of three points generated using CD40L^WT^, CD40L^A208D^, and CD40L^K143A^ with the degree of correlation quantified by the Pearson’s r-coefficient. (E) Correlation maps (assessed by Pearson’s r coefficient for all readouts as shown in C, D and Figure S7) showing data of CD40 signaling and downstream functions measured in Farage and PBMC B cells (indicated) using the indicated experiments (from left to right: MTP, phosphorylation of p38 and Erk1/2 by TGT, and phosphorylation of p38 and Erk1/2, TFEB translocation, and Ig class-switch by PMFA) correlate poorly with the 2D affinity (*A*_c_*K*_a,0_) and reciprocal off-rate (1⁄*K*_off,0_) measured in the absence of force (0 pN, top half), yet strongly with the same binding parameters (*A*_c_*K*_a_(15pN), 〈*t*_*b*_〉(15pN)) when measured at the optimal force (15 pN, bottom half).

After extensive correlative analyses, we found that, among all binding parameters tested, *A*_c_*K*_a_(15pN) and 〈*t*_*b*_〉(15pN), two force-dependent binding parameters evaluated at the optimal force where the CD40–CD40L^WT^ catch bond transitions to the slip bond, stand out as the best correlators for CD40-mediated B cell signaling and function (Figures 7C-D and S7). This can be clearly seen from the Pearson’s r correlation maps showing much better correlations of *A*_c_*K*_a_(15pN) and 〈*t*_*b*_〉(15pN) than *A*_c_*K*_a,0_ and 1⁄*k*_off,0_ with all the results of the MTP, TGT, and PMFA experiments (Figure 7E), highlighting a key role of force in regulating function through modulating binding.

## Discussion

The CD40–CD40L interaction is a critical element for the communication between B and T cells, and hence orchestrates antibody-mediated immune responses.^64^ Effective T-B communication in GC ensures the production of high-affinity, class-switched antibodies to provide appropriate and long-lasting immunity against pathogens. Dysregulation of the CD40– CD40L interaction causes immune deficiencies, such as the X-HIgM, characterized by abnormal antibody isotype production and increased susceptibility to opportunistic infections. Surprisingly and unlike other types of receptor–ligand interactions, nothing is known about the mechanobiology of the CD40–CD40L system and whether mechanotransduction plays a role in X-linked Hyper IgM syndromes. Here, we have uncovered the crucial role of mechanotransduction in regulating CD40 signal output that impacts B cell signaling and function. We also showed that X-HIgM-associated variants of CD40L were unable to induce proper CD40 mechanotransduction, which is likely to contribute to the antibody deficiency syndromes in affected patients. These results were achieved by using a suite of biomechanical techniques, some of which have been well established in our lab,^32,39,40^ while others were newly developed, and never have all been integrated and used collectively to examine the mechanotransduction of a single immunoreceptor as was done in this study.

The first technique is BFP that analyzes mechanically receptor–ligand interaction kinetics at the single-bond level. We found that compared to CD40L^WT^, X-HIgM CD40L mutants display a reduced zero-force 2D affinity for CD40. Analysis of bond dissociation under force showed that CD40 forms strong catch-slip bonds with CD40L^WT^. While this is quite interesting on its own right, it is even more striking to find that CD40 forms weakened catch-slip bonds or even slip-only bonds with X-HIgM CD40L mutants (Figures 1I-J and S1). The utility and usefulness of these measurements have been demonstrated by their power of predicting B cell signaling and function (Figure 7C-E and S7).

The second technique is MTP that measures B cell internal forces on CD40. Again, the observation that B cells exert endogenous forces on CD40 is by itself interesting, but it is even more intriguing to find that B cells tune the endogenous force that they exert on CD40 to match the level sustainable by the CD40 anchor as demonstrated by three lines of evidence (Figures 2 and S2). First, when CD40 was anchored to the surface *via* WT CD40L, the forces B cells exerted were between 12 and 19 pN, which closely matches the optimal force where the lifetime of the CD40–CD40L^WT^ catch-slip bond peaks as it transitions from the catch regime to the slip regime. Second, when CD40 was anchored to the surface *via* an anti-CD40 antibody, the number of CD40 experiencing endogenous forces decreases percipetually as the level of force increased, matching the slip-only bond profile of the CD40–anti-CD40 bonds. Third, when the CD40 was anchored to the surface *via* X-HIgM CD40L mutants, the ability for B cells to exert forces on CD40 was greatly reduced, matching the suppressed catch-slip bond and the slip-only bond profiles of these interactions. It should be pointed out that how the MTP force readout depends on the binding parameters is not fully understood. To generate a force signal, CD40 has to form durable bonds with MTP-tagged surface-bound CD40L or anti-CD40 antibodies capable of sustaining force, hence depending on the binding parameters. However, this alone is insufficient because beads coated with soluble CD40 ectodomain can bind CD40L with the same binding parameters as B cell surface CD40 but would not generate any force signal. Therefore, binding-induced signaling must be required to activate the force generation and transmission machinery inside the B cell to result in a force signal in the MTP experiment. Our results suggest that B cells mechanically sample CD40L by exerting endogenous forces on CD40.

The third technique is TGT that caps endogenous forces on CD40. Not only did we employ TGT to test cell spreading as previously done by others,^44,47,65^ but we also extended the previous coverslip-based technique (Figure 3A) to a bead-based method (Figure 3D) amenable for analysis of intracellular signaling, enabling us to find that endogenous forces are important to proper B cell spreading and enhance CD40 signaling (Figures 3 and S3). Indeed, when the endogenous forces allowed to apply on CD40–CD40L bonds were capped by <12 pN, B cells could not efficiently spread on CD40L and the phosphorylation of relevant signaling molecules was reduced compared to the case when bonds were allowed to experience forces up to 56 pN. In contrast to CD40L^WT^, X-HIgM mutations to CD40L failed to support appreciable spreading and abrogated signaling regardless of whether forces were restricted to <12 pN or allowed to be up to 56 pN (Figures 3 and S3). Similar to MTP, how the TGT force readout depends on the binding parameters is not fully understood. Different from MTP, which reports the number of CD40– CD40L bonds on which an above threshold force is applied by the B cell, TGT reports the force requirement on CD40–CD40L bonds for function. Together, these two techniques provide different and complementary perspectives of B cell endogenous forces induced by and exerted on CD40–CD40L bonds and the functional consequences of not letting the B cell exert such forces.

The fourth technique is PMFA, which turns the table around to examine the effects of exogenous instead of endogenous forces. Three types of CD40-mediated B cell functions were quantified: intracellular signaling (short-term), TFEB translocation (mid-term), and Ig class switch (long-term). Compared to the TGT experiment finding that limiting endogenous force decreases signaling (Figure 3E-H), the PMFA experiment found that exerting exogenous force increases signaling (Figure 4D-G), showing two sides of the same coin. X-HIgM mutants, instead, are defective in their ability to induce force-dependent enhancement of CD40 signaling (Figures 4D-G and S4G-I), also consistent with the MTP results (Figures 2H-I and S2). Analogously to proximal CD40 signaling, externally applied force on CD40–CD40L bonds induced prolonged nuclear localization of TFEB, necessary for the induction of secondary signals to rescue B cells from BCR-induced apoptosis, while X-HIgM mutations to CD40L abrogate this prolonged nuclear localization of TFEB (Figures 5D, E and S5). Enabled by our PMFA *in vitro* CSR assay, we demonstrated that force on CD40–CD40L bonds promote antibody class-switch in GC and memory B cells as well as B cells as a whole. While in the absence of force X-HIgM CD40L mutants show no difference in their ability to induce CSR compared to CD40L^WT^, in the presence of force these mutants downregulate CSR (Figures 6B-E and S6). Thus, CD40 mechanotransduction enables B cells to distinguish between CD40L^WT^ and its X-HIgM mutants in terms of their ability to induce Ig class-switch recombination as CD40 could not discriminate between them without applied force. In other words, for the CD40 mutants to manifest defects in X-linked Hyper IgM syndrome, their participation in Ig class-switch recombination must involves CD40 mechanotransduction.

The remarkable ability of CD40–CD40L interaction kinetics to be regulated by force, manifesting as catch-slip bond formation demonstrated in this study, adds to a growing list of immunoreceptor–ligand interactions that have been observed to form catch-slip bonds, highlighting the importance of mechanotransduction through immunorecepters.^27^ In contrast to integrins whose catch bonds are mostly linked to their adhesive functions,^66–73^ the CD40–CD40L catch bond has been shown to impact signaling and function. Previously, the connections of TCR,^39^ pre-TCR^74^, chimeric antigen receptor,^75^ Fcγ receptor IIA,^76^ and NKG2D’s^77^ catch bonds to signaling and function of T cells, neutrophils, and NK cells have been established by applying external forces to thesereceptor–ligand bonds while observing intracellular calcium fluxes concurrently^39,78^ and by correlating altered catch bonds and functions *via* mutations in the receptors and/or ligands.^39,74–82^ In the present work, by comparison, the immunological relevance of the CD40–CD40L catch-slip bond has been more thoroughly demonstrated. We employed four powerful approaches to obtain convincing evidence: 1) matching the optimal force of the catch-slip bond to the optimal force that B cells generate, 2) suppressing B cell signaling by capping endogenous forces below the optimal level, 3) applying external forces to CD40–CD40L bonds while monitoring B cell signaling and function concurrently, and 4) correlating altered catch bonds, signaling and function *via* CD40L mutations found in X-HIgM patients (Figures 7 and S7). This would not have been possible without the technical innovation in the TGT and PMFA assays and without the organic integration of the four powerful mechanobiological techniques discussed earlier.

Whereas CD40 signaling is decreased by capping endogenous force and increased by applying exogenous force, differences in the effect size and timing have been observed between the TGT (Figures 3E-H and S3) and PMFA (Figures 4D-G and S4) experiments. Such differences can be reconciled by considering the size difference between magnetic beads used in the TGT and PMFA beads, as well as the different CD40L densities used in the two experiments. The magnetic beads are unlikely able to support endogenous force generation due to their smaller size and the lower CD40L density required to limit the number of formed bonds per bead-cell contact to be ∼2. In sharp contrast, the much larger TGT beads functionalized with much higher CD40L density would allow their formation of a much greater number of CD40 bonds with B cells over a much larger contact area, which should promote endogenous force generation. Moreover, it is possible that while endogenous forces may rise and decay after contact formation, leading to a similar rise and fall of CD40 signaling in the TGT experiment, in the PMFA experiment bonds between CD40 and CD40L-coated magnetic beads were subject to continuous external force, favoring sustained p38 and Erk1/2 phosphorylation.

Interestingly, the phosphorylation of p38 and Erk1/2 are differentially enhanced by force, much greater for p38 than Erk1/2. This indicates that CD40 mechanotransduction may impact some signaling pathways more than others. Differential responses to mechanotransduction have been observed in other systems. For example, mechanosensing through platelet glycoprotein Ib switches integrin α_IIb_β_3_ from the low-affinity state to the intermediate affinity state, but does not do so for integrin α_V_β_3_.^73^ The mechanism of this differential regulation of signaling by CD40 mechanotransduction needs to be addressed in future studies.

The physiological sources of external forces applied to CD40 are of great interest. As mentioned earlier, the mechanical interaction between highly mobile T and B cells predicts that CD40–CD40L bonds are likely subjected to physical forces.^27^ In addition, due to the ability of T cells to generate endogenous force on their membrane receptors, such as TCR,^40,45,83^ Program Death 1 (PD-1),^46^ and Lymphocyte Function-Associated antigen 1 (LFA-1),^65^ it is conceivable that T_FH_ cells could also apply endogenous force to CD40–CD40L bonds to balance the endogenous B cell force and thus promote the elevated and sustained CD40 signaling, TFEB translocation, GC reaction, and Ig class-switch. Whether T_FH_ cells do indeed generate the force on CD40L and how such force may also affect CD40L signaling is an open question worthy of future exploration.

To conclude, our findings establish that mechanical forces regulate the CD40–CD40L interaction and shed light on the mechanistic underpinnings of X-HIgM syndrome by demonstrating that dysregulation of mechanotransduction mediated by CD40–CD40L bonds underlies the pathology observed in X-HIgM patients. This study improves our understanding of T-B cell interactions and potentially opens new therapeutic avenues for the treatment of X-HIgM.

## Acknowledgments

This work was supported by NIH grants U01CA250040 (C.Z.) and U01CA280984 (A.S. and C.Z.) and by a grant from the Hyper IgM Foundation (C.Z.). H.-K.C. was partly supported by a National Research Foundation grant of South Korea (2021R1A6A3A03038382).

## Author contributions

Conceptualization, H.-K.C., S.T., C.Z., A.S., and J.W.; Methodology, H.-K.C., S.T., M.M., R.G., Z.Z., J.L., D.M.R.-A., C.Z., J.W., and A.S.; Investigation, H.-K.C., S.T.; Visualization, H.-K.C. and S.T.; Writing – Original Draft, H.-K.C., S.T., and C.Z.; Writing – Review & Editing H.-K.C., S.T., M.M., J.W., A.S., and C.Z.; Funding Acquisition, H.-K.C., J.W., A.S., and, C.Z.; Project administration, C.Z.

## RESOURCE AVAILABILITY

### Lead contact

Further information and requests for resources and reagents should be directed to the corresponding author, Cheng Zhu.

### Materials availability

Unique reagents generated in this study (*e.g*., DNA and protein sequencies, plasmids and strains) are available from the corresponding author upon request.

### Data and code availability

Data presented in the main body and supplemental information plus codes used in this study are presented in (https://github.com/Chengzhulab/Mechanobiology-of-CD40). Other analysis procedures are described in the methods, as well as the quantification and statistical analysis sections.

## EXPERIMENTAL MODEL AND SUBJECT DETAILS

### Cell lines

To generate the protein producer cell line HEK293T/17 BirA^+^, HEK293T/17 cells (purchased from ATCC) were transduced with the plasmid pHR-CMV-TetO2_HA-BirA-ER encoding a green fluorescence protein (GFP) marker and HA-tagged ER-resident *E. coli* biotin ligase (a gift from A. Radu Aricescu, Addgene plasmid # 113897) to allow for *in vivo* biotinylation at the N-terminus of CD40L ectodomain. HEK293T/17 BirA^+^ cells were subsequently transduced with soluble CD40L-encoding lentiviruses to generate producer cell lines stably expressing each soluble CD40L construct. HEK293T/17 BirA^+^ CD40L^+^ cells were then sorted with their GFP marker to equalize expression. Farage B lymphoma cells were a kind gift from Wendy Béguelin (Weill Cornell Medical College) and maintained in RPMI 1640 supplemented with 10% FBS, 1% P/S and 10 mM HEPES. Ramos cells (from Jürgen Wienands’lab), were maintained in RPMI 1640 supplemented with 10% FBS, 1% P/S, 10mM HEPES, 1 mM L-glutamine, 1 mM sodium pyruvate, and 50 µM β-mercaptoethanol. PBMC derived B cells were purified from freshly isolated blood of healthy donors according to the protocol approved by the Institute Review Broad of the Georgia Institute of Technology. Briefly, whole lymphocytes were separated by centrifugation with Histopaque®-1077 (Millipore Sigma) from which CD19^+^ B cells were purified using the EasySep™ Human CD19 Positive Selection Kit II (STEMCELL Technologies). After purification, PBMC B cells were kept in RPMI supplemented with 10% FBS, 1% P/S, 10mM HEPES, and 1X NEAA, and used immediateily for each experiment.

## METHOD DETAILS

### Generation of soluble CD40L constructs

Soluble WT CD40L (CD154) gene (TNF homology region M113-L261) including a N-terminal 6xHis tag and AviTag was synthesized and purchased from Twist Bioscience and subcloned into the pHR-CMV-TetO2_3C-Avi-His6 IRES-EmGFP vector (a gift from A. Radu Aricescu, Addgene plasmid # 113888) by in vivo assembly (IVA) at the AgeI and XhoI sites. Mutations to the WT CD40L gene were obtained by site-directed mutagenesis using primers purchased from IDT technologies and KLD enzyme mix purchased from New England Biolabs (Ipswich, MA). The sequence of the cloned genes was verified by whole-plasmid nanopore sequencing from Plasmidsaurus and Sanger sequencing of insert from Eurofins Genomics. Lentiviruses encoding for all CD40L constructs were generated by PEI transfection of HEK293T/17 cells (ATCC) with the CD40L transfer vectors as well as packaging and envelope plasmids, psPAX2 and pMD2.G (gifts from from Didier Trono, Addgene plasmids # 12260 and # 12259). Lentiviral particles were harvested from the supernatant and concentrated 100X using PEG precipitation, using the Lenti-X concentrator (Takara Bio). Stable expression of soluble CD40L constructs was generated by lentiviral transduction of HEK293T/17 cells expressing BirA enzyme for *in vivo* biotinylation. His-tagged CD40L proteins were harvested from cell supernatant and purified by Ni-NTA chromatography. After purification, soluble proteins were buffer exchanged and concentrated using Amicon® Ultra-15 10K Centrifugal Filter units (MilliporeSigma) and stored in PBS + 30% glycerol for further use after snap freezing in liquid nitrogen. Purified proteins were then verified by SDS-PAGE as well as by capturing proteins on streptavidin beads and immunocytometry with anti-CD40L and anti-His tag antibodies (Biolegend), and compared to commercial soluble CD40L (ACROBiosystems).

### BFP measurement

BFP setup: Our custom-designed and home-built BFP has been previously described,^37,84,85^ and was used in this study to measure the 2D kinetics of bonds between a panel of soluble CD40L constructs or anti-CD40 antibody and either soluble and membrane-bound CD40 in the absence of force by the adhesion frequency assay^32,34^ and in the presence of force by the force-clamp assay.^39,42^ In brief, we coated streptavidin-conjugated glass beads with soluble CD40L constructs or anti-CD40 antibody using biotin-streptavidin coupling. We then attached a functionalized bead to the apex of a biotinylated RBC aspirated on a glass micropipette to form a force-transducer (probe), which has millisecond (time), nanometer (position), and piconewton (force) resolutions.^37,85^ Such a force probe was used to test binding with CD40 coated on beads or expressed on lymphoma or PBMC B cells (targets) in repeated mechanical cycles (Figures 1B,C and S1C). All measurements were conducted in the chamber buffer (L15 with 1% BSA and 5 mM HEPES) at room temperature.

Adhesion frequency assay: During each mechanical cycle, we drove the target aspirated on a glass micropipette using a piezotranslator controlled by a computer program to approach and briefly (0.1-5 s) contact with a small impingement force (∼ −20 pN) to allow bond formation, followed by retraction of the target at a force loading rate of 1000 pN/s. This generates a binary readout: either a binding event manifesting a positive force (tension) upon retraction, or a no binding event manifesting the return of negative force (compression) to zero as the target moves away from the probe (Figure S1C, lower right). We estimated the adhesion frequency (*P*_a_ = # binding events divided by total # of contacts) from repeated mechanical cycles of the same contact time (*t*_c_) using the same probe-target pair, converted *P*_a_ to average number of bonds per contact by a logmarith transformation ⟨*n*⟩ = − ln(1 − *P*_a_), divided the value by the site densities of CD40 (*m*_CD40_) and CD40L (*m*_CD40L_) measured by independent flow cytometry experiment, obtained Mean ± SEM values using 3-4 probe-target pairs, measured these for a range of contact times, and fitted the data to a previously published model:^32^ 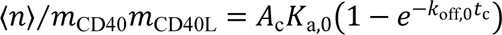 to evaluate the effective 2D affinity *A*_c_*K*_a,0_ (in μm^4^) and off-rate *K*_off,0_ (in s^−1^), where *A*_c_ (in μm^2^) is the contact area whose exact value is difficult to determine precisely but was ket constant for each probe-target system. For this reason, we lump it together with *K*_a,0_ and add an adjective “effective” to distinguish *A*_c_*K*_a,0_ from the 2D affinity*K*_a,0_ (in μm^2^). For the same reason, caution must be exercised in comparing *A*_c_*K*_a,0_ values measured from different systems, *e.g*., bead-bead (cf. Figure 1B) and bead-cell (cf. Figure 1C) experiments or bead-cell experiments with different cells, even for the same molecular interactions because the contact areas may not be the same. Also note that the subscript 0 associated with the two binding parameters indicate that the *K*_a,0_ and *K*_off,0_ values are measured under zero-force conditions.^32^

Force-clamp assay: In the retraction phase of the mechanical cycle, if a bond survived the ramping and reached a preset tension level, the force was clamped until spontaneous bond dissociation and a pair of values of the clamped force *f* and bond lifetime *t*_b_ were measured (Figure S1C, upper right). To ensure that most adhesion events were mediated by single molecular bonds, we controlled the adhesion to be infrequent (≤ 20%).^32^ Bond lifetimes were measured at forces ranging from 2 – 40 pN, pooled, and binned into >5 force bins (>35 measurements per bin) to reduce system errors, and presented as mean bond lifetime 〈*t*_b_〉 ± SEM. The SEM of mean force is usually smaller than the size of the symbols in the plots, which is obscured.

### MTP surface preparation

We functionalized DNA-based MTP on glass coverslips using a protocol adapted from previous publications.^45,46^ In brief, 25 mm glass coverslips were sonicated in 50% ethanol for 15 min and rinsed 6X with di-H_2_O. Following rinsing, the coverslips were etched in 40 ml Piranha solution (2:1 ratio of sulfuric acid to H_2_O_2_) for 30 min. Coverslips were washed 6X with di-H_2_O and 3X with 100% ethanol. The surfaces were then silanized with 3% APTES in ethanol for 1 h, then washed 3X in ethanol and dried at 80C for 30 min. After drying, 200 μl of 10 mg/ml LA-PEG, 50 mg/ml mPEG in 0.1 M NaHCO_3_ was added to each surface and incubated overnight at 4C°. Surfaces were then washed 3X with di-H_2_O and 200 μl of 1 mg/ml Sulfo-NHS-Acetate was added to two coverslips placed together as a sandwich and incubated for 30 min at room temperature. Coverslips were again rinsed 3X with di-H_2_O and 500 μl Au-NP solution was added to each sandwich and incubated for 1 h at room temperature. During Au-NP incubation, 300 nM hairpin strand, 330 nM quencher strand, and 330 nM Atto647 strand in 1 M NaCl were annealed in a thermocycler by heating to 95 °C for 5 min and gradually cooling (−5 °C/min) to 25 °C. After rinsing Au-NP coated surfaces with 3X di-H_2_O and 2X 1 M NaCl washes, 100 μl of DNA probes were added to each sandwich and incubated overnight at 4 °C. The following day, the MTP coverslips were rinsed with 3X PBS washes and 40 μg/ml streptavidin in PBS was added to each sandwich and incubated for 1 h at room temperatue. Following 3X rinse with PBS, 40 μg/ml of CD40L construct or anti-CD40 antibody in PBS + 2% BSA was added to each sandwich and incubated for 1 h at room temperature.

### TGT surface preparation

To prepare TGT surfaces, 25×75 mm^2^ coverslips were sonicated, etched, and silanized as described above for MTP and then dried under argon stream after 3X ethanol washes. After drying, one Ibidi Sticky-Slide VI 0.4 was mounted on each coverslip to create 6 microfluidic channels per coverslip while making sure to remove any bubbles from the adhesive areas. NHS-PEG4-Azide was then diluted in 0.1M NaHCO_3_ to 10 mg/ml and 50 μl of the solution added to each channel and incubated at room temperature for 1 h. The channels were then washed 3X with 1 mL di-H_2_O and blocked with PBS + 1% BSA for 1 h. During this incubation, the DBCO-bottom strand and the biotin-top strand of TGT probes (220 nM each in 1 M NaCl) were annealed in a thermocycler by heating to 95 °C for 5 min and gradually cooling (−5 °C/min) to 25 °C. After blocking, the channels were washed 3X with PBS and 50 μl of PBS was left in the channels to prevent drying. The TGT probes were then added (50 μl) to each channel and incubated overnight at room temperature to covalently functionalize the channels with TGT probes *via* strain-promoted alkyne-azide cycloaddition (SPAAC). The following day, channels were rinsed 3X with PBS, and 50 μl of 20 μg/ml streptavidin solution in PBS were added to each channel and incubated for 1 h at room temperature. After rinsing 3X with PBS, 50 μl of 20 μg/ml biotinylated CD40L in PBS + 1% BSA was added to each channel and incubated for 1 h at room temperature.

### TGT bead preparation

TGT probes were heat annealed as described above (220 nM in 1M NaCl for each strand, 1 mL in a thermocycler. After annealing, the required amount of azide-coated PMMA beads (12 μm, PolyAn) for a 1:1 cell:bead ratio were washed 2X with PBS + 2% BSA by centrifugation at 200 g for 3 min, then resuspended in the TGT probe solution (∼1 mL) and incubated overnight on rotor at room temperature to allow functionalization of the beads with the TGT probes by SPAAC click reaction. The following day, TGT-coated beads were washed 2X with PBS + 2% BSA and resuspended in 500 μl of 40 μg/ml streptavidin in PBS + 2% BSA and incubated for 1 h on rotor at room temperature. After washing 2X with PBS + 2% BSA, beads were resuspended in 1 μg/ml (for Farage B cells) or 3 μg/ml (for PBMC B cells) of biotinylated CD40L in PBS + 2% BSA and incubated for 1 h at room temperature. TGT beads were then washed 2X in PBS + 2% BSA to remove free CD40L before use.

### Parallel magnetic force activation (PMFA) assay

To apply magnetic force in a parallel fashion across all wells of a 24-well plate, we designed, in Solidworks, a lid that would fit the top of a 24-well plate with 2 slots per well to house magnets and 3D printed it out of Polylactic acid (PLA) filament. Using such a lid, we mounted two 5 mm cube neodymium magnets in antiparallel configuration per well, making sure that all magnets on the lid are mounted following the exact same antiparallel pattern to allow reproducible force application in all wells. The magnetic lid could then be sterilized by immersion in 70% EtOH and then dried under argon stream. Next, to the center of each well of an non-tissue culture treated 24-well plate, we applied a droplet of 5-min epoxy using a pipette tip. At the same time, 5 mm-diameter glass coverslips were cleaned by sequential washes in 70% EtOH and di-H_2_O. Using a pair of tweezers, each coverslip was then dried by gentle tapping of its side on a kimwipe, and gently placed on top of the epoxy droplet in the center of the well. To aid this process, it is helpful to mark the center of each well on the back side of the well plate, and to proceed by applying epoxy and adding coverslips row-by-row, as waiting too long will result in the epoxy hardening and the inability to attach the coverslips to the plate. After attaching coverslips to the center of every well, the wells were washed with 1 ml PBS, then PBS was added again to each well (∼1 mL) and the plate was incubated for 30 min at room temperature. Following incubation, PBS was thoroughly aspirated out of each well, and then plates were left to air dry completely for 30 min at room temperature. After wells and coverslips were fully dried, an 18 μl droplet of 0.01% PLL solution (Sigma-Aldrich) was carefully added to each coverslip, and plates were incubated at 4 °C overnight to allow coating of PLL on the glass coverslips. The following day, wells were washed with ∼1 ml PBS and dried by aspiration, making sure that the area around and underneath the coverslips is fully dried. After drying, 18 μl of B cells in R10 (1 – 2 × 10^5^ cells/18 μl) were added on top of each coverslip and were incubated for 10 min at 37 °C to allow cells to settle and become immobilized on the PLL-coated coverslips. After immobilization, 2 μl of a well-mixed solution of CD40L-coated magnetic beads in R10 was carefully added on top of the R10 droplet above the immobilized cells, at the center of every well. Plates were then incubated for 10 min at 37 °C to allow the beads to settle on the cells by gravity and to start interacting with B cells. After beads were settled, 80 μl of R10 (with supplements or treatments depending on the specific assay) was added to each well around the coverslip. Critically, care must be taken while adding the 80 μl of R10 to each well, such that the coverslips and cells are not touched and that the flat droplet of media does not contact the side walls of the well, which would result in meniscus formation and unwanted drying of the sample. We find that adding 80 μl of media slowly while making circular motions around the coverslip leads to best results. After incubation for 10 min at 37 °C, the magnetic lid was gently placed on top of the plates (or a normal lid without magnets for control samples), making sure that the magnets do not directly contact the top of the media droplet, and cells were incubated under force application for a given time depending on the specific experiment.

### PMFA for intracellular signaling

For this experiment, PMFA plates, cells and beads were prepared as described above, adding 2×10^5^ cells per well. Beads were functionalized with 1 μg/ml CD40L for Farage cells or 3 μg/ml CD40L for PBMC B cells. In the samples treated with soluble CD40L, an equivalent amount of CD40L was added to each well as that of CD40L coated on beads (2 μl of 1 or 3 μg/ml for Farage or PBMC B cells, respectively). Following preset incubation times (0, 0.5, 1, 3, or 5 h) cells were harvested by trypsinization (500 μl TrypLE, Gibco) for 5 min at 37 °C, followed by 1 ml of R10. Cells were then washed with PBS and fixed in 1 ml of 1X PBS with 4% PFA for 10 min at 37 °C. After fixation, cells were permeabilized by adding the cells to 9 ml of ice-cold 100% methanol, to a final concentration of 90% methanol and incubated on ice for 30 min. Cells were then washed twice in 1 ml FACS buffer and resuspended in 50-100 μl of 1:50 diluted primary PE-conjugated antibodies against phosphorylated p38 or Erk1/2 (Cell Signaling Technology) and stained for 1 h at room temperature on a rotor. After staining, cells were washed twice in 1 ml FACS buffer and resuspended in 0.2-0.3 ml FACS buffer for flow cytometer analysis.

### PMFA for TFEB translocation

In this experiment, PMFA plates and magnetic beads were prepared as described above. Ramos cells (2.5×10^5^ cells/well) were added to each coverslip with the addition of 8 μg/ml anti-IgM for BCR stimulation. Beads coated with 0.2 μg/ml CD40L were then added to each well as described above, and cells were incubated under magnetic force application for 1 h. After incubation, cells were harvested by trypsinization as described above, and 4 wells were combined for each sample to reach 1×10^6^ cells/sample. After washing with 500 μl PBS by centrifuging at 300 g for 5 min, cells were fixed with 500 μl of 1:1 mixture of PBS and CytoFix buffer (BD) for 20 min at room temperature. After fixing, cells were washed with 1 ml of FACS buffer by centrifuging at 700 g for 5 min at 4 °C and then permeabilized and stained in 150 μl of FACS buffer + 0.1% Triton X-100 with primary anti-TFEB antibody (1:150 dilution) for 30 min at 4 °C in the dark. After washing twice with 1 ml FACS buffer by centrifuging at 700g for 5 min at 4 °C, cells were stained with secondary anti-rabbit FITC antibody in FACS buffer (1:150 dilution) for 30 min at °C in the dark. Cells were then washed twice in 1 ml FACS buffer by centrifuging at 700 g for min at 4 °C and resuspended in 30 μl of FACS buffer containing 10ug/ml 7-AAD for nuclear staining. Cells were then imaged with the ImageStream cytometer in the brightfield, FITC, and 7-AAD channels, at 60X magnification and gated for in-focus and live cells.

### PMFA for class-switch

In this experiment, we stimulated PBMC B cells using the PMFA setup described above. Specifically, we immobilized 10^5^ PBMC B cells on PLL coated coverslips in an 18 μl R10 droplet and added 2 μl of magnetic beads coated with CD40L (0.2-0.6 μg/ml coating) to each well at a 1:2 cell to bead ratio (2×10^5^ beads per well). After addition of the beads, 80 μl of R10 supplemented with 100 ng/ml BAFF and 20 ng/ml IL-4 (final concentration) was added around the coverslips to form a 100 μl droplet as described above. Following a brief incubation for 10 minutes at 37 °C, magnetic lids were placed above the plates containing samples to be stimulated under magnetic force, and cells were incubated for 6 h under force. After this 6-h incubation, magnetic lids were removed, 400 μl of R10 supplemented with 100 ng/ml BAFF and 20 ng/ml IL-4 was added to each well (500 μl total volume), and all cells and beads were transferred to new 24 well plates for prolonged stimulation without force to elicit germinal center formation.

After 4 days, plates were centrifuged at 450 g for 5 min, and 250 μl of the conditioned media was removed from the top of each well, making sure cells and beads were not aspirated. We then added 500 μl of R10 supplemented with 100 ng/ml BAFF and 10 ng/ml IL-21 to each well and further incubated cells and beads for 4 days at 37 °C. On day 8, cells were harvested and washed twice with 200 μl FACS buffer after transferring to a 96-well plate by centrifuging at 500 g for 5 min. After washing, dead cells were stained with 100 μl of LIVE/DEAD Fixable stain (ThermoFisher) at 1000X dilution in PBS for 30 min at room temperature in the dark with shaking. Cells were then washed twice in 200 μl FACS buffer by centrifuging at 500g for 5 min and resuspended in 100 μl antibody mix solution in FACS buffer, with anti-CD27, anti-CD38, anti-IgM, anti-IgG (all 1:200 dilution) and anti-CD19 (1:100 dilution). Cells were thus stained for 1 h at room temperature in the dark with shaking. After staining, cells were washed twice with 200 μl FACS buffer by centrifuging at 500 g for 5 min and then fixed in 100 μl of 1X PBS with 4% PFA for 30 min at room temperature in the dark with shaking. Following fixation, PFA was removed by two washes with 200 μl FACS buffer by centrifuging at 600 g for 5 min, and cells were then resuspended in 200 μl FACS buffer and stored at 4 °C in the dark until flow cytometry analysis. Flow cytometry gating was determined based on fluorescence minus one (FMO) control for each fluorophore.

## QUNATIFICATION AND STATISTICAL ANALYSIS

Briefly, custom LabView and MATLAB softwares were written for analysis of BFP and MTP data. Data obtained from TGT and PMFA assays were analyzed using FlowJo and MATLAB softwares. Statistical details have also been provided in figure legends where applicable. For additional details, see the STAR METHODS section.

### Designing magnetic force profile in PMFAA

To design the configuration of magnet-lid and chamber to generate a well-defined magnetic force on magnetic beads in a well-defined space, a well-defined magnetic field gradient is needed. Positive magnetophoresis is a phenomenon that occurs when magnetic objects (*e.g.*, magnetic beads) possess positive magnetization vectors and are immersed in non-magnetic substances such as buffer, cells, or tissues. In this context, a relative magnetic permeability 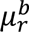 (≡ *μ*_*b*_/*μ*_0_ where *μ*_*b*_ and *μ*_0_ are the permeability values of the magnetic particle and of vacuum, respectively), is always higher than that of buffer, cells, or tissues (which is iessentially non-magnetic (*i.e*., *μ*_*m*_ ≈ *μ*_0_)). In the presence of a gradient in the external magnetic field, these magnetic objects tend to be attracted to spaces with denser magnetic fields and higher field gradients. For a spherical magnetic particle, the magnetic force is given by,

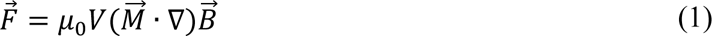

Where 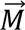 is the magnetization of a particle per unit volume, *V* is the volume of the magnetic particle, and 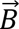 is the applied external magnetic field in the surrounding media. Since we used a superparamagnetic bead that has minimal residual magnetization, the magnetic force can be exerted on the beads only when there is an external magnetic field. If the magnetic field itself is strong enough, by Langevin’s theory of paramagnetism, the magnetization vector reaches a saturated value of 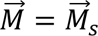, the saturated magnetization per unit volume. Hence, the magnetic force can be reduced to simple form such that 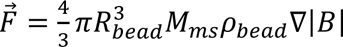 where *ρ*_bead_ is the density of a magnetic bead (1.4 g/cm^3^ for M270), *R*_*bead*_ is the radius of a magnetic bead (1.4 μm for M270), and *M*_*ms*_ is the mass saturation magnetization (10.88 J/T/kg for M270).^86–88^ To calculate the magnetic field, in the absence of both electric fields and external currents, the magneto-static state for hard ferromagnets (two permanent neodymium magnets with a magnetization grade of N50) with a 1 mm gap in between can be described as,

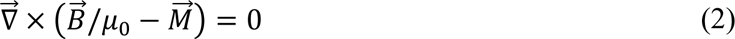

Where 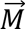 is the magnetization from the remanent field (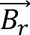/*μ*_0_ for *B*_*r*_ = 1.24 T for a pair of two N50-grade magnets)^89^. Using the above equations, due to the finite size constraint of cell media in each well of a 24 well-plates, we calculated the magnetic force profile under the circumstance where each pair of magnets is placed as low as possible (1.5 mm from the surface) inside the wells.

### Analysis of image flow cytometer

All images obtained from the Amnis ImageStream were exported to Matlab and converted to Tiff format for further analysis of TFEB translocation. The raw data format consists of one cell per image, and includes 4 channels, namely brightfield, side scatter, FITC (for TFEB) and 7-AAD (for nucleus). First, the nucleus was segmented based on the 7-AAD channel by applying adaptive threshold to binarize the raw image. Similarly, each entire cell was segmented based on the FITC channel. After segmenting both whole cell and nucleus, the mean background fluorescence was subtracted from each TFEB and 7-AAD image. To calculate the similarity index between the TFEB channel and the nucleus channel, we computed the Pearson’s coefficient (*p*) between the two channels for the whole-cell segmented area (thus excluding the background pixels from the calculation). The Pearson’s coefficient between two images *x* and *y* is defined as:

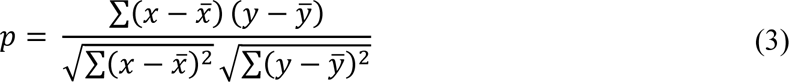

where 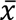 and 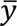 are the mean pixel values of each image. The Pearson’s coefficient is a dimensionless value between −1 and 1 representing the linear correlation between the pixel values of the two images, where 1, −1, and 0 indicate perfect correlation, perfect inverse correlation, and no correlation, respectively. To address the low dynamic range of the Pearson’s coefficient, another metric has been defined as a monotonic function of the Pearson’s coefficient, the similarity score (S):

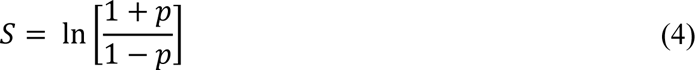

The similarity score effectively rescales the Pearson’s coefficient from −∞ to +∞, allowing for more accurate measurements of similarity between the two channels. Next, to determine the portion of cells with nuclear TFEB, a gate was applied to the similarity score distribution, with a threshold set at 1. Thus, cells with a similarity score > 1 were identified as having nuclear TFEB, while cells with a score < 1 were classified as having cytosolic TFEB.

