## Supplemental Information for "Mechanotransduction governs CD40 function and underlies X-linked Hyper IgM syndrome"

### Supplemental Figures

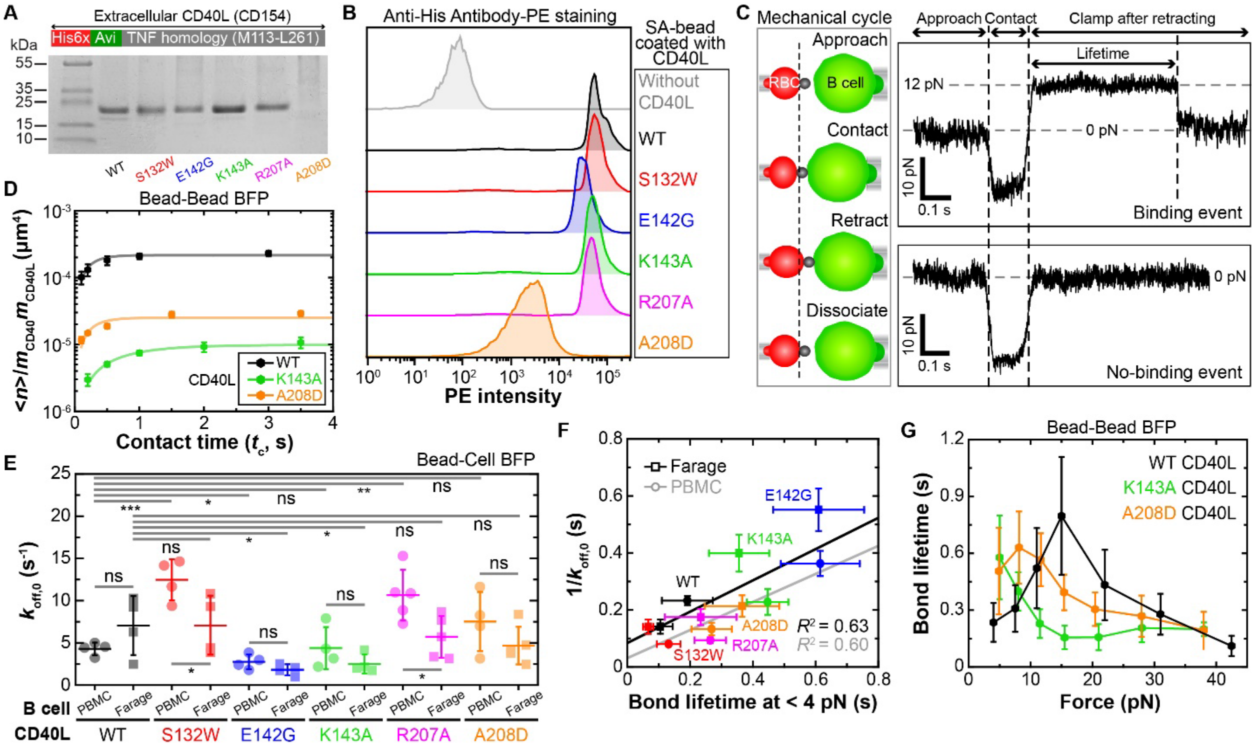


**Figure S1. BFP measurements of CD40 interactions with CD40L constructs, Related to Figure 1**

(A) Domain diagram (upper) and representative SDS-PAGE gel image (lower) of soluble CD40L constructs (indicated on the bottom of each lane from WT to point-mutations related to X-HIgM type I). The first lane of the gel indicates molecular mass. A 6xHis tag and an Avi-tag are added to the N-terminus for detection and coating on beads (below).

(B) Representative flow-cytometric analysis of indicated soluble CD40L constructs coated on glass beads *via* streptavidin-biotin coupling and detected by PE-conjugated anti-His antibody.

(C) Schematics of the mechanical cycle used in BFP measurements (left box) and representative force *vs* time traces in two approach-contact-retraction-dissociation cycles of the force-clamp assay that resulted in a binding event with a lifetime measured under force-clamp mode (upper right) and a no-binding event (lower right).

(D) Mean ± SEM ($n$ = 4) of average number of bonds per contact ($\langle n \rangle$) normalized by site densities of CD40 ($m_{\mathrm{CD40}}$) and CD40L ($m_{\mathrm{CD40L}}$) *vs* contact time ($t_{\mathrm{c}}$) plots measured from the bead-bead BFP experiment (cf. Figure 1B). $\langle n \rangle$ $(= -\ln(1 - P_{\mathrm{a}}))$ was converted from the adhesion frequency ($P_{\mathrm{a}}$ = # binding events divided by total # of contacts) directly measured using the indicated (by color) CD40L constructs coated on probe beads interacting with soluble CD40 ectodomain coated on the target beads. Data (points) were fitted (curves) by a previously published model:[1] $\langle n \rangle / m_{\mathrm{CD40}} m_{\mathrm{CD40L}} = A_{\mathrm{c}} K_{\mathrm{a},0}(1 - e^{-k_{\mathrm{off},0} t_{\mathrm{c}}})$. Thus, the steady-state levels of the curves reflect the effective 2D affinities of CD40 for the different CD40L constructs.

(E) Mean ± SD of zero-force off-rate ($k_{\mathrm{off},\,0}$) measured by the bead-cell BFP experiment (cf. Figure 1C) using PBMC (circles) and Farage (squares) B cells to interact with the indicated (by color) CD40L constructs. Scatters represent values measured from individual bead-cell pairs ($n$ = 4-5). Two-sided t-test was used to assess significance (ns > 0.05, $0.01 < * < 0.05$, $0.001 < ** < 0.01$, and $0.0001 < *** < 0.001$).

(F) Plot of reciprocal zero-force off-rate ($1/k_{\mathrm{off},\,0}$) measured from the adhesion frequency assay *vs* Mean ± SEM of bond lifetime $\langle t_b \rangle (< 4\mathrm{pN})$ measured at the lowest force bin by the force-clamp assay using the indicated (by color) CD40L constructs to interact with CD40 from Farage (squares) or PBMC (circles) B cells. Data were fitted by a straight line for each system and the goodness-of-fit is indicated by the $R^2$ value.

(G) Mean ± SEM of bond lifetime *vs* force plots ($n$ > 35 lifetime measurements per force bin) of beads bearing indicated CD40L constructs interacting with beads coated with CD40 ectodomain measured using the force-clamp mode shown in C (bead-bead BFP), showing a strong catch-slip bond for $\mathrm{CD40L^{WT}}$, suppressed catch-slip bond for $\mathrm{CD40L^{A208D}}$, and slip-only bond for $\mathrm{CD40L^{K143A}}$, consistent with the bead-cell BFP results shown in Figure 1I-J.

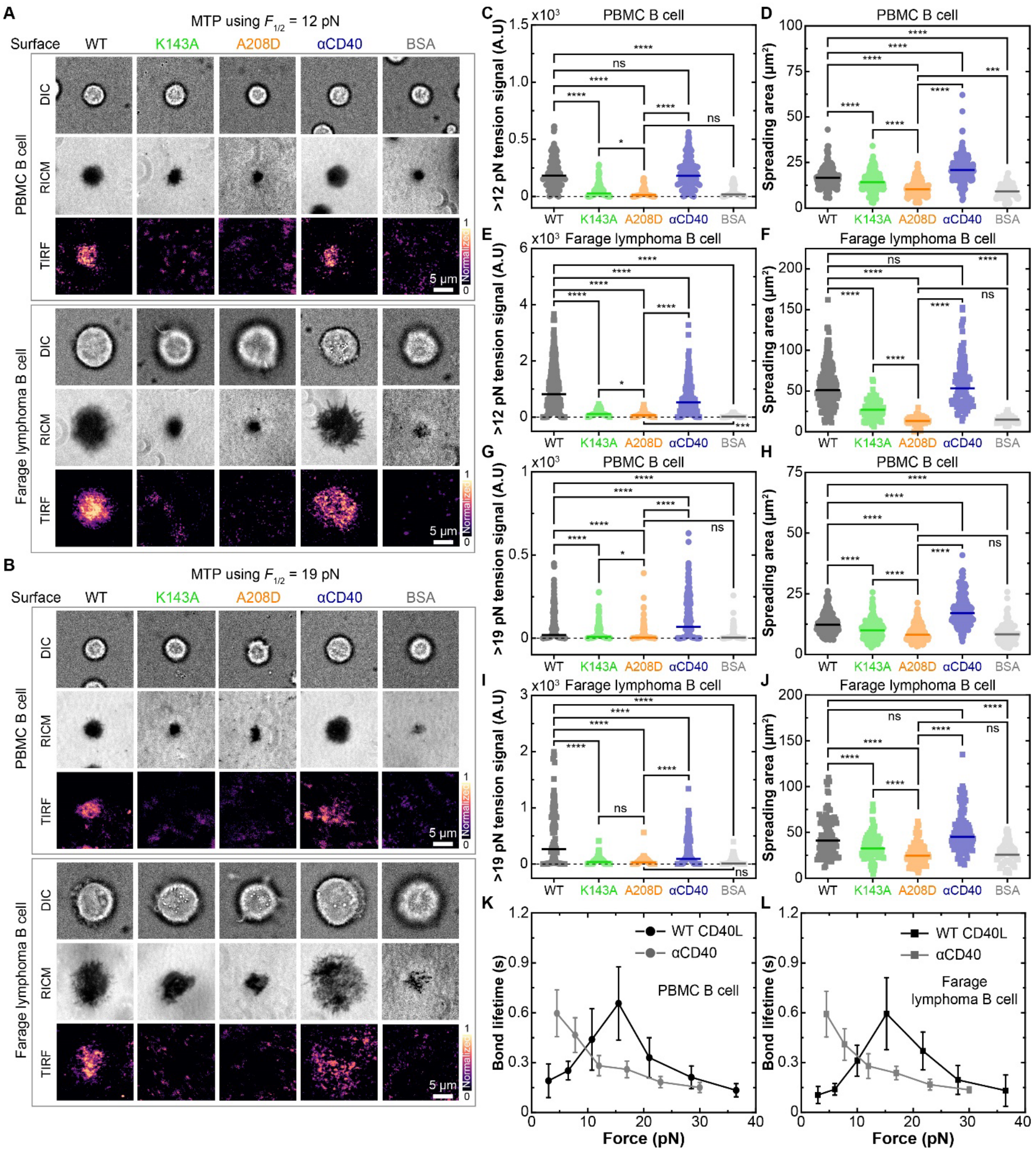


### Figure S2. Molecular tension probe experiments with 12- and 19-pN threshold forces, Related to Figure 2

(A and B) Representative images of MTP experiments. PBMC (upper box) and Farage (lower box) B cells were placed on MTPs of 12 pN (A) or 19 pN (B) threshold force tagged with the indicated CD40L constructs, anti-CD40 antibody, or BSA, imaged by differential interference contrast microscopy (DIC, 1st row), reflection interference contrast microscopy (RICM, 2nd row), and total internal reflection fluorescence normalized tension signal (TIRF, 3rd row). Scale bar = 5 μm.

(C-J) Mean and individual data points (*n* = 39-262) of tension signals (C, E, G and I) and spreading areas (D, F, H and J) of PBMC (C, D, G and H) and Farage (E, F, I and J) B cells on MTPs with 12 pN (C-F) or 19 pN (G-J) threshold force functionalized with indicated CD40L constructs, anti-CD40 antibody, or BSA. Two-sided t-test was used to assess significance of differences between groups (ns > 0.05, 0.01 < ∗ < 0.05, 0.001 < ∗∗ < 0.01, 0.0001 < ∗∗∗ < 0.001, and ∗∗∗∗ <0.0001).

(K and L) Mean ± SEM of bond lifetime *vs* force plots (*n* > 35 lifetime measurements per force bin) of anti-CD40 antibody (new data) or WT CD40L (replotted from Figure 1I-J for comparison) interacting with CD40-expressing PBMC (K) or Farage (L) B cells measured by bead-cell BFP (cf. Figure 1C) force-clamp experiments (cf. Figure S1C).

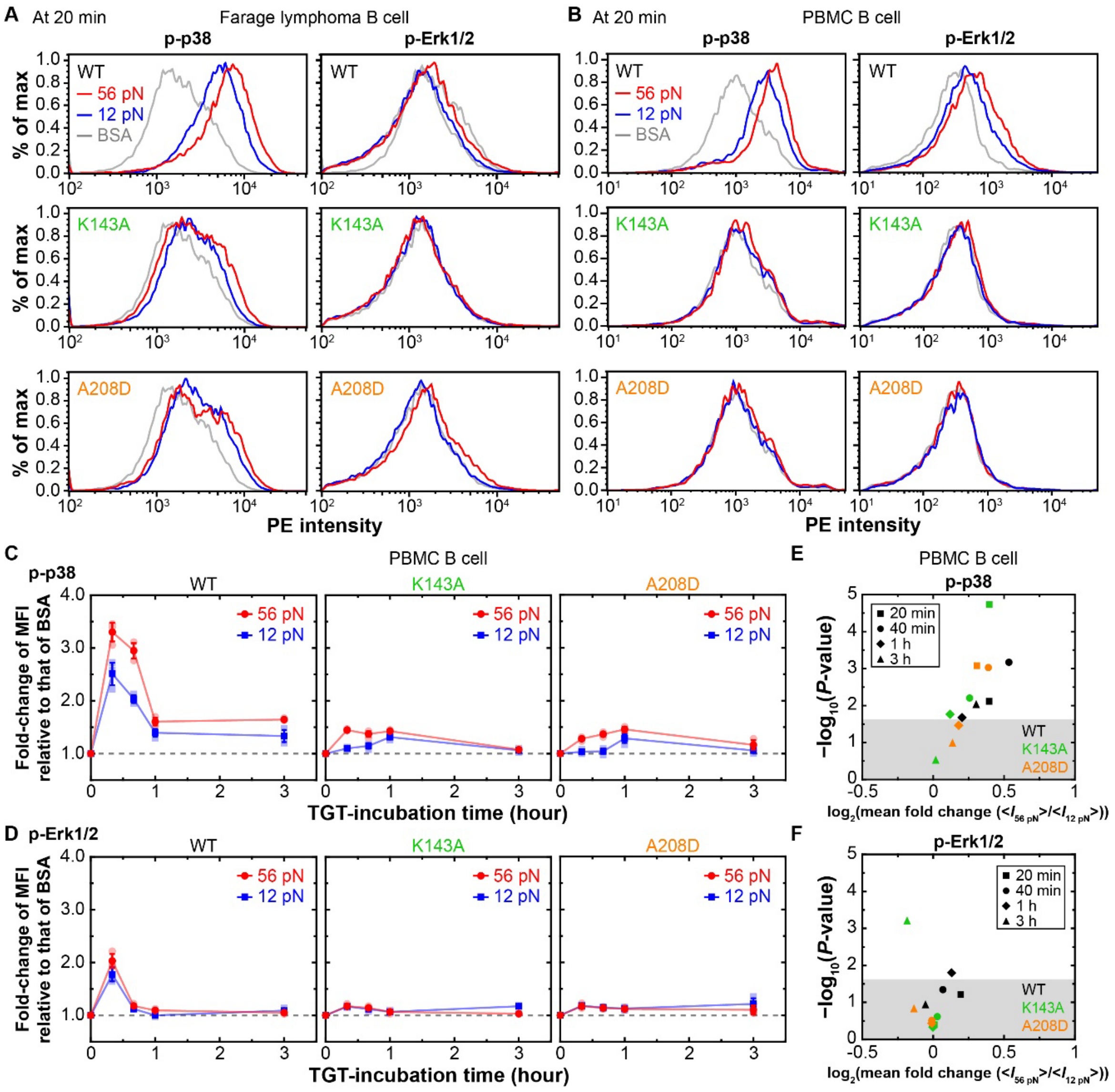


**Figure S3. Time-course of CD40 signaling under limited endogenous forces by TGT, Related to Figure 3**

(A and B) Representative flow cytometry data showing intracellular staining of PE-conjugated antibody against phospho-p38 (p-p38) (left) and phospho-Erk1/2 (p-Erk1/2) (right) induced by incubating Farage (A) or PBMC (B) B cells with TGT beads of 12 pN (blue) or 56 pN (red) threshold force tagged with BSA control (gray) or CD40L$^{WT}$ (top), CD40L$^{K143A}$ (middle), or CD40L$^{A208D}$ (bottom), for 20 min.

(C and D) Mean ± SD with individual data points ($N = 3$ with $n > 10{,}000$ cells) of fold-change of mean fluorescence intensity (MFI) of PE-conjugated antibody (normalized by that of BSA) against p-p38 (C) and p-Erk1/2 (D) in PBMC B cells stimulated with TGT beads of 12 pN (blue) or 56 pN (red) threshold force tagged with $CD40L^{WT}$ (left), $CD40L^{K143A}$ (middle), or $CD40L^{A208D}$ (right) are plotted *vs* cell-bead incubation time.

(E and F) $Log_{10} - Log_2$ plots of relative change of *P*-value *vs* mean fold change of MFI of p-p38 (E) and p-Erk1/2 (F) in PBMC B cells stimulated by TGT beads when the threshold force were increased from 12 to 56 pN. Different points represent different TGT-incubation times and different colors indicate different CD40L constructs. Shaded area indicates *P*-value > 0.05 above which represents the region of statistical significance.

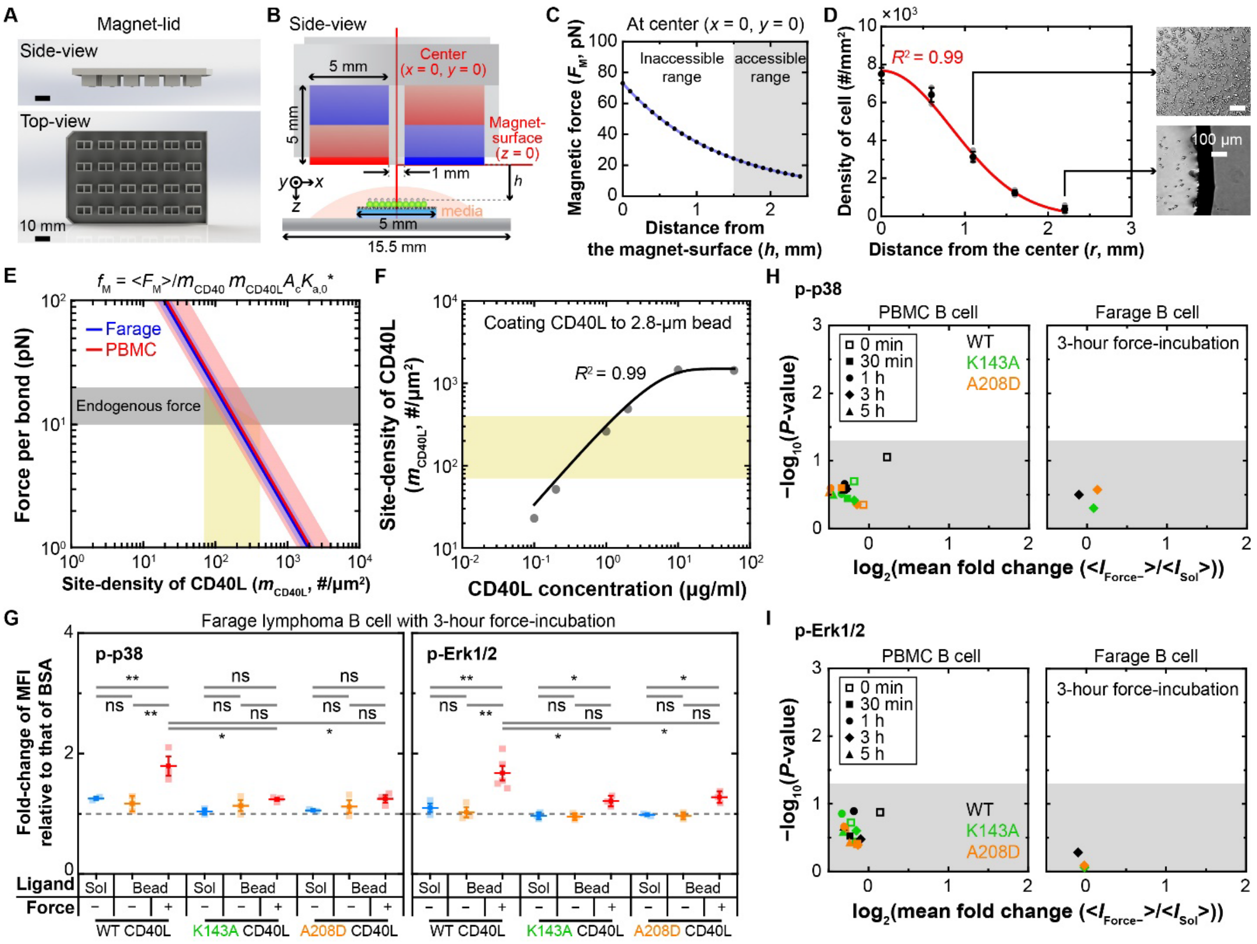


**Figure S4. Parallel magnetic force activation (PMFA) assay design and application, Related to Figures 4**

(A) Schematic drawing (upper: side view, lower: top view) of a magnetic lid designed by *Solidworks* software. The magnetic lid is fitted to a commercial 24 well plate. Scale bar = 10 mm.

(B) Schematic of PMFA from the side view with detailed dimensions. Center is defined as the middle line in the gap between a pair of magnets (vertical red line). The distance from the magnet surface is defined as the height of magnet from the sample.

(C) Calculated magnetic force with respect to height along the center line ($(x, y, z) = (0,0, h)$) defined in B.[2,3] Shaded area indicates the experimentally accessible region. The curve is obtained from experimentally driven double-exponential fitting with respect to the height of magnets.

(D) Mean ± SD with individual data (N = 3) of cell density along radius of coverslip after seeding cells in each plate well. The red curve represents a Gaussian fit with the goodness-of-fit indicated by the $R^2$. Insets are representative images at the corresponding locations. Scale bar = 100 μm.

(E) Initial force per bond *vs* site density of CD40L on the beads calculated using force-free 2D affinity. The colored lines and corresponding shaded areas are determined from the respective BFP results of Farage (blue) and PBMC (red) B cells. The gray shaded area indicates a target range of force corresponding to the endogenous force measured in the MTP assay. The yellow box shows appropriate site densities of CD40L on the bead for the PMFA assay. Of note, the number of bonds for a single bead decay with time because bonds would dissociate over time under external force and the beads would be pulled towards the magnets, preventing rebinding to the cells.

(F) Optimization of CD40L site density *vs* CD40L concentration during coating. Points are measured using flow cytometry. Curve represents trendline. The yellow box shows appropriate range of CD40L site densities on the bead for the PMFA assay.

(G) Relative fold-change of MFI (normalized by MFI from the BSA control) of p-p38 (left) and p-Erk1/2 (right) in Farage cells after 3-hour incubation with soluble or magnetic bead-coated CD40L constructs (indicated) with or without force applied by using the PMFA assay. Two-sided t-test was used to assess significance of differences between groups (ns > 0.05, $0.01 < * < 0.05$, $0.001 < ** < 0.01$, and $0.0001 < *** < 0.001$).

(H and I) $Log_{10}$ – $Log_2$ plots of relative change of *P*-value *vs* mean fold change of MFI of PE-conjugated antibody against p-p38 (J) or p-Erk1/2 (K) in PBMC (left) and Farage (right) B cells stimulated by CD40L constructs (indicated by color) when the molecular presentation was changed from soluble form to bead-bound form without force application. Different scatter points represent different incubation times. Shaded area indicates *P*-value > 0.05 above which represent the region of statistical significance.

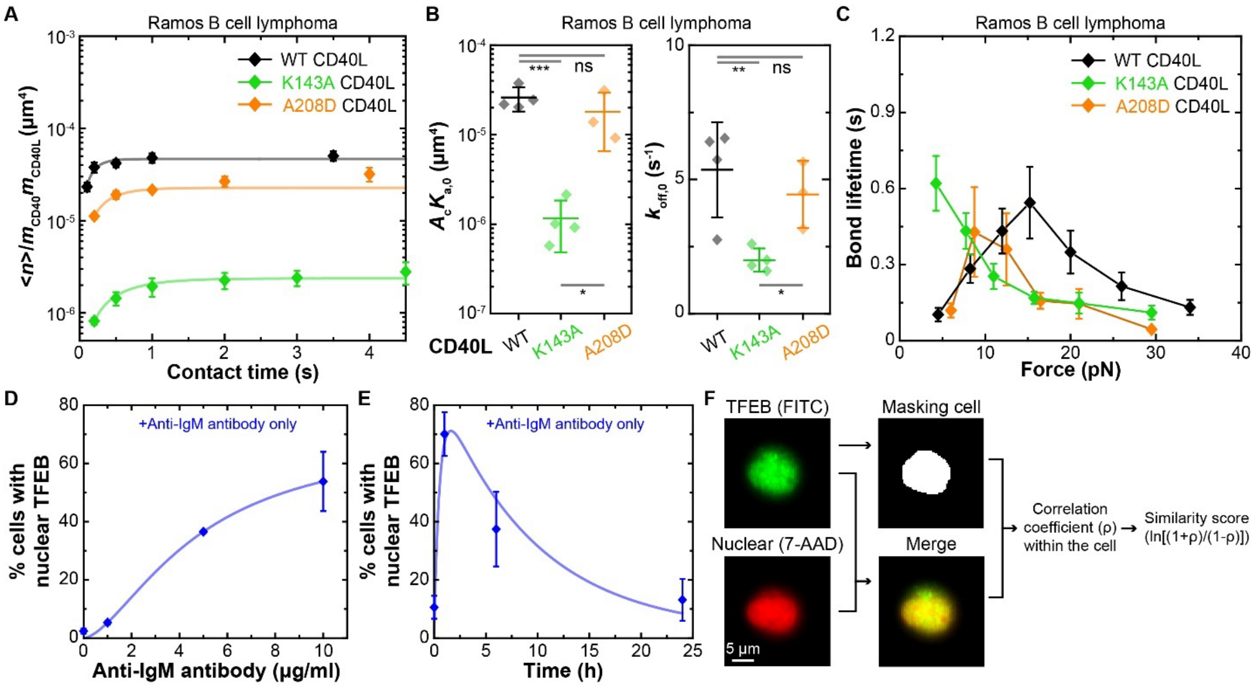


**Figure S5. Kinetic measurements of CD40–CD40L interactions on Ramos B cells and time and dose response curves of TFEB translocation, Related to Figures 5**

(A) Plots of Mean ± SEM ($n$ = 3-4) of average number of bonds per contact ($\langle n \rangle$) normalized by the site densities of CD40 ($m_{\mathrm{CD40}}$) and CD40L ($m_{\mathrm{CD40L}}$) *vs* contact time ($t_{\mathrm{c}}$). $\langle n \rangle$ ($= -\ln(1-P_{\mathrm{a}})$) was converted from the adhesion frequency ($P_{\mathrm{a}}$ = # binding events divided by total # of contacts) directly measured with the indicated (by color) CD40L constructs coated on probe beads interacting with CD40-expressing Ramos B cells by the adhesion frequency assay using the bead-cell BFP setup (cf. Figure 1C). Data (points) were fitted (curves) by a previously published model:[1] $\langle n \rangle / m_{\mathrm{CD40}} m_{\mathrm{CD40L}} = A_{\mathrm{c}} K_{\mathrm{a,0}} \left(1 - e^{-k_{\mathrm{off,0}} t_{\mathrm{c}}}\right)$. Thus, the steady-state levels of the curves reflect the effective 2D affinities of CD40 for the different CD40L constructs.

(B) Mean ± SD of zero-force effective 2D affinity ($A_{\mathrm{c}} K_{\mathrm{a,0}}$) (left) and off-rate ($k_{\mathrm{off,\,0}}$) (right) evaluated from the curve fits in A. Scatters indicate data measured using individual bead-cell pairs ($n$ = 3-4). Two-sided t-test was used to assess statistical significance among different CD40L constructs (ns > 0.05, $0.01 < * < 0.05$, $0.001 < ** < 0.01$, and $0.0001 < *** < 0.001$).

(C) Mean ± SEM of bond lifetime *vs* force plots ($n > 35$ lifetime measurements per force bin) of beads coated with indicated CD40L constructs interacting with CD40-expressing Ramos B cells measured using the bead-cell BFP (cf. Figure 1C) force-clamp assay (cf. Figure S1C).

(D and E) % cells with nuclear TFEB *vs* concentration of anti-IgM antibody (D) and incubation time (E) used to activate TFEB translocation replotted from a previous study.[4] Lines are fitting curves from (D) simplified Hill-equation: $E_{\max}/(1 + (EC_{50}/[\alpha\text{IgM}])^n))$ and (E) simple rise-fall-kinetic equation of signaling: $A\left(e^{-k_2 t} - e^{-k_1 t}\right)/(k_1 - k_2)$ where $E_{\max}$, $EC_{50}$, $n$, $A$, $k_1$ and $k_2$ are fitting parameters. Using this information, the amount of anti-IgM antibody and incubation time were selected for the present study.

(F) Strategy employed to analyze similarity scores. Cell can be masked to isolate whole-cell and nuclear areas from FITC (TFEB) and 7-AAD (nucleus) channels. Based on the 2-dimensional Pearson's correlation coefficient of the FITC and 7-AAD signals within the segmented cellular area, we calculated a similarity score between the two channels (See Quantification and Statistical Analysis in METHODS).

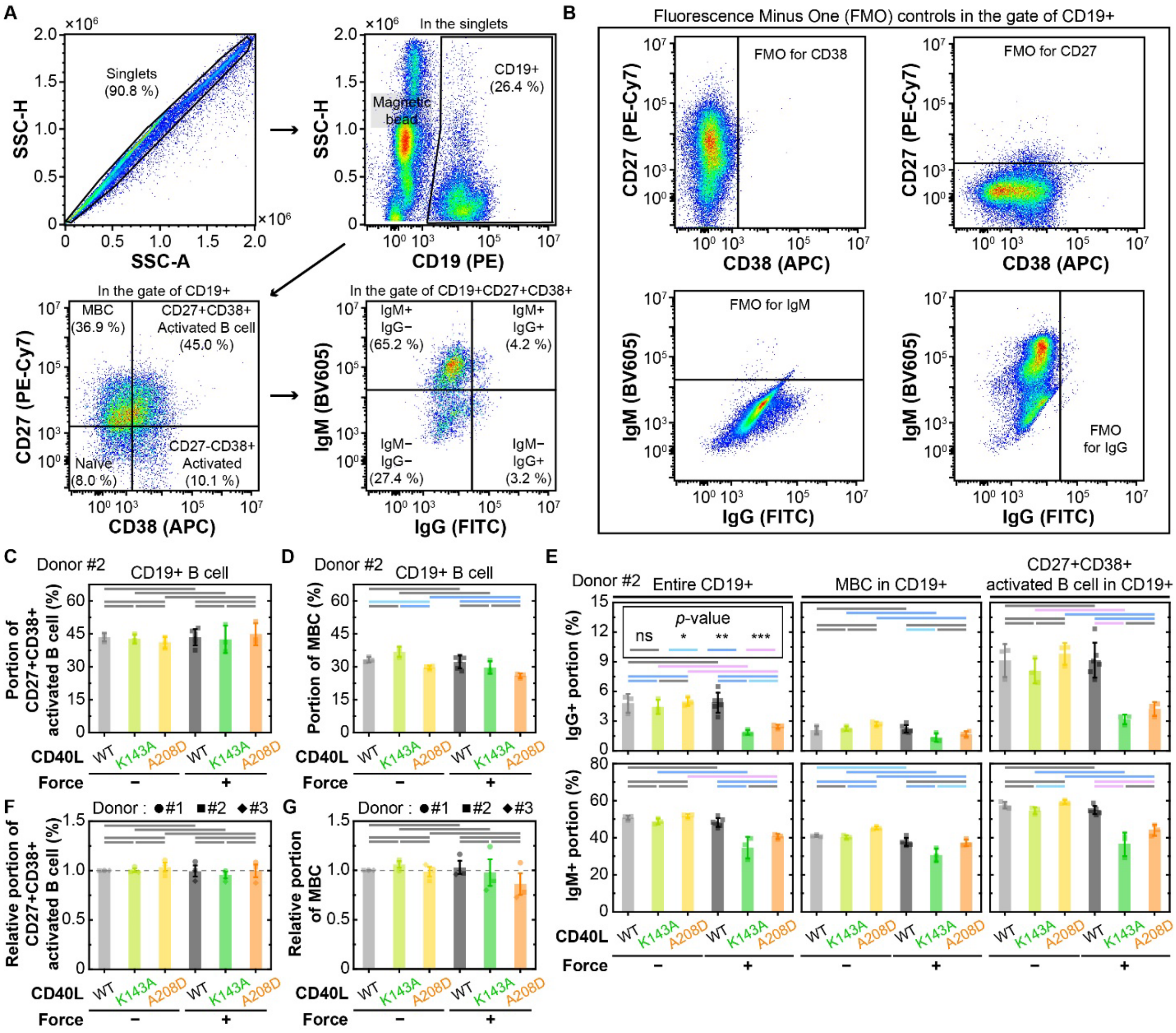


**Figure S6. Ig class switch recombination experiment combined with PMFA assay, Related to Figure 6**

(A) Gating strategy of flow cytometry data to measure Ig class switch recombination (arrows between panels indicate the gating sequence). The first gating is from singlets in SSC-H *vs* SSC-A domain (upper left). A second gating can be done in CD19 (PE) *vs* SSC-H domain (upper right). Among CD19$^+$ cells, a third gating can be set in CD38 (APC) *vs* CD27 (PE-Cy7) domain (lower left). Lastly, Ig class switch recombination can be checked in IgG (FITC) *vs* IgM (BV605) domain (lower right).

(B) Strategy for setting gates by Fluorescence Minus One (FMO) controls. FMO is used to find the minimum bound of staining intensity, providing the standard for threshold level of positive signal. Each plot shows indicative FMO examples.

(C and D) Mean ± SD with individual data points ($N = 3$ with $n > 10{,}000$ cells) of $CD19^{+}CD27^{+}CD38^{+}$ activated B cell subpopulation (C) and Memory B cell (MBC) subpopulation (D) among the entire population of donor #2 $CD19^{+}$ B cells stimulated with magnetic beads bearing indicated CD40L constructs with or without the magnet lid in the PMFA assay.

(E) Mean ± SD, and individual data points, of $IgG^{+}$ (top row) and $IgM^{+}$ (bottom row) portions in all $CD19^{+}$ B cells ($1^{st}$ column), MBC ($2^{nd}$ column), and $CD19^{+}CD27^{+}CD38^{+}$ activated B cells ($3^{rd}$ column) from donor #2.

(F and G) Mean ± SEM (data from 3 different donors) of fold-change of $CD19^{+}$ $CD19^{+}CD27^{+}CD38^{+}$ activated B (F) and MBC (G) cell subpopulations after stimulation of PBMCs by the indicated CD40L constructs coated on magnetic beads pulled by force in the PMFA experiments, normalized by the data of WT CD40L stimulation without force.

Two-sided t-test was used to assess statistical significance among different CD40L constructs in C-F (ns > 0.05, $0.01 < * < 0.05$, $0.001 < ** < 0.01$, and $0.0001 < *** < 0.001$).

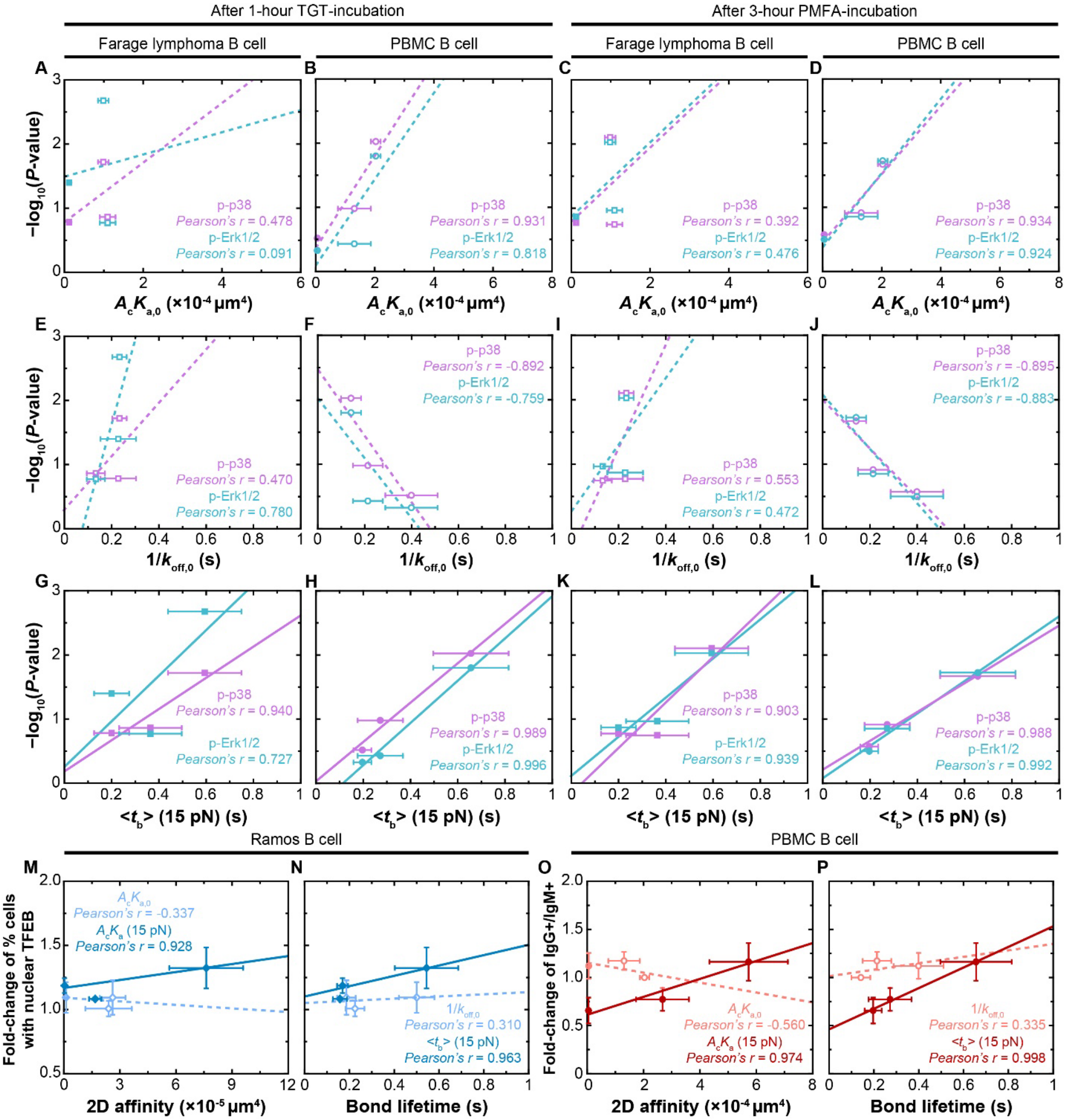


**Figure S7. Correlation plots between B cell functional readouts and 2D binding parameters in the presence or absence of force.**

(A-L) The *P*-values (assessed using two-sided t-test) quantifying the statistical significance in the differences between groups of signaling (evaluated by p-p38 and pErk1/2 levels) in Farage (A, C, E, G, I, and K) or PBMC (B, D, F, H, J, and L) B cells stimulated by three CD40L constructs functionalized on TGT beads of 12- or 56-pN force threshold for 1 h (A, B, E, F, G, and H, reproduced from Figures 3G,H and S3E,F) or magnetic beads with or without force application by PMFA for 3 h (C, D, I, J, K, and L, reproduced from Figures 4F,G) are plotted *vs* zero-force effective 2D affinity $A_{\mathrm{c}}K_{\mathrm{a,0}}$ (A-D), zero-force reciprocal off-rate $1/k_{\mathrm{off,0}}$ (E-F, I-J) and bond lifetime at 15 pN $\langle t_b \rangle(15\mathrm{pN})$ (G-H, K-L) reproduced from Figures 1H and S1E (for $A_{\mathrm{c}}K_{\mathrm{a,0}}$ and $k_{\mathrm{off,0}}$) or from Figure 1I-J (for $\langle t_b \rangle(15\mathrm{pN})$). Data (points) are presented as Mean ± SEM (*n* = 3-4 bead-cell pairs for A-H and ~35 measurements for I-L) and fitted by a straight line for each group of three points generated using CD40L$^{\mathrm{WT}}$, CD40L$^{\mathrm{A208D}}$, and CD40L$^{\mathrm{K143A}}$ with the degree of correlation quantified by the Pearson's r-coefficient.

(M-P) Fold-change of % cells with nuclear TFEB in Ramos B cells (M and N, reproduced here from Figure 5E) or ratios of IgG$^{+}$/IgM$^{+}$ of GCB cells (O and P, reproduced from CD19$^{+}$CD27$^{+}$CD38$^{+}$ activated B cells of Figure 6E) stimulated by three CD40L constructs with and without force by PMFA were plotted *vs* 2D affinities at zero-force ($A_{\mathrm{c}}K_{\mathrm{a,0}}$, light blue in M and light red in O) or 15-pN force ($A_{\mathrm{c}}K_{\mathrm{a}}(15\mathrm{pN})$, dark blue in M and dark red in O) and *vs* zero-force reciprocal off-rate ($1/k_{\mathrm{off,0}}$, light blue in N and light red in P) or bond lifetime at 15 pN ($\langle t_b \rangle(15\mathrm{pN})$, dark blue in N and dark red in P) reproduced from previous Figures. Data (points) are presented as Mean ± SEM (*N* = 3 for the *y* values and *n* = 3-4 bead-cell pairs for zero-force parameters and ~35 measurements for parameters at 15 pN for the *x* values) and fitted by a straight line for each group of three points generated using CD40L$^{\mathrm{WT}}$, CD40L$^{\mathrm{A208D}}$, and CD40L$^{\mathrm{K143A}}$ with the degree of correlation quantified by the Pearson's r-coefficient.
